# Genomic analysis based on chromosome-level genome assembly reveals an expansion of terpene biosynthesis of *Azadirachta indica*

**DOI:** 10.1101/2021.11.11.468207

**Authors:** Yuhui Du, Wei Song, Zhiqiu Yin, Shengbo Wu, Jiaheng Liu, Ning Wang, Hua Jin, Jianjun Qiao, Yi-Xin Huo

## Abstract

*Azadirachta indica* (neem), an evergreen tree of the Meliaceae family, is a source of the potent biopesticide azadirachtin. The lack of a chromosome-level assembly impedes the understanding of in-depth genomic architecture and the comparative genomic analysis of *A. indica*. Here, a high-quality genome assembly of *A. indica* was constructed using a combination of data from Illumina, PacBio, and Hi-C technology, which is the first chromosome-scale genome assembly of *A. indica*. The genome size of *A. indica* is 281 Mb anchored to 14 chromosomes (contig N50=6 Mb and scaffold N50=19 Mb). The genome assembly contained 115 Mb repetitive elements and 25,767 protein-coding genes. Evolutional analysis revealed that *A. indica* didn’t experience any whole-genome duplication (WGD) event after the core eudicot γ event, but some genes and genome segment might undergo recent duplications. The secondary metabolite clusters, TPS genes, and CYP genes were also identified. Comparative genomic analysis revealed that most of the *A. indica*-specific TPS genes and CYP genes were located on the terpene-related clusters on chromosome 13. It is suggested that chromosome 13 may play an important role in the specific terpene biosynthesis of *A. indica*. And the gene duplication events may be responsible for the terpene biosynthesis expansion in *A. indica*. This will shed light on terpene biosynthesis in *A. indica* and facilitate comparative genomic research of the family Meliaceae.

## 1. Introduction

*Azadirachta indica* (neem) is a member of Meliaceae family, which is extensively studied for its bioactive products(Schmutterer, 1995). It grows natively on the Indian subcontinent and also in other countries such as Egypt and the Kingdom of Saudi Arabia. *A. indica* is a source of abundant limonoids and simple terpenoids which are responsible for its biological activity(Dai, Yaylayan, Raghavan, Pare, & Liu, 2001). Azadirachtin, the most important activate compound in the neem tree, has been intensively studied because of its wide range of insecticidal profile and low toxicity to mammalian(Ley, 1994). Additionally, the neem tree extracts also exhibit many pharmaceutical functions, such as anti-inflammatory, anticancer, antimicrobial, and antidiabetic activities(Abdelhady, Bader, Shaheen, El-Malah, & Barghash, 2015; Soares et al., 2014). A lot of studies have focused on the synthesis of azadirachtin, including chemosynthesis, hairy root culture, cell line culture, and callus culture(Mithilesh & Rakhi, 2014; Rodrigues, Festucci-Buselli, Silva, & Otoni, 2014; Srivastava & Srivastava, 2013; Veitch et al., 2007). However, these methods are either of low extraction efficiency or not environmentally friendly. Therefore, the reconstruction of biosynthetic pathway of azadirachtin for heterologous production is an alternative method.

Omics strategy is an effective method to study the biosynthesis of secondary metabolite. Transcriptomes of *A. indica* tissues (stem, leaf, flower, root, and fruit) have been sequenced, which paved the way to the potential synthetic pathway of azadirachtin and gene expression profiles in various organs. Draft genomes have also been sequenced, which led to a basic understanding of the genetic characteristics of *A. indica*(Krishnan, Jain, Gupta, Hariharan, & Panda, 2016; Krishnan et al., 2012; Kuravadi et al., 2015). However, the lack of a chromosome-level genome sequence has hindered fully understanding of the secondary metabolite biosynthesis and the evolution of *A. indica*. In addition, Meliaceae are known to produce around 1,500 structurally diverse limonoids, which have agricultural and medical values(Hodgson et al., 2019). A chromosome-level genome is essential for genome-wide studies of the Meliaceae family.

In this study, the first chromosome-level genome of *A. indica* was assembled through a combination of Illumina, PacBio, and Hi-C technology. Based on the assembled genome sequence, annotation, evolutionary analysis, whole-genome duplication (WGD), secondary metabolite clusters identification, and resistance gene identification were performed. These results improved the understanding of the genomic architecture of *A. indica*. This chromosome-level genome assembly can be used as a new reference genome for *A. indica*, laying a substantial foundation for further genomic studies.

## 2. Materials and Methods

### 2.1 Plant material, DNA preparation, and genome sequencing

Fresh tissues of *A. indica* were randomly collected from the park of Hainan University (100.61438 E, 36.28672 N), Hainan Province, China. Fresh leaves were collected to isolate genomic DNA of *A. indica* for de novo sequencing and assembly. For Illumina sequencing, a paired-end library (270 bp) was generated and sequenced on the Illumina HiSeq X Ten platform. For PacBio sequencing, a 20 kb insert library was generated and sequenced on the PacBio RSII platform.

### 2.2 Genome assembly

First, Canu v2.0(Koren et al., 2017) software was used to correct and assemble raw PacBio sequencing reads, and 886 contigs with N50 ~ 6M were assembled by Canu. In addition, we performed a round of polishing on the assembled contigs using the RACON(Vaser, Sovic, Nagarajan, & Sikic, 2017) with the PacBio long reads, and the polished contigs were further corrected two rounds on the genome-wide base-level by Pilon v1.21 (Walker et al., 2014) with the Illumina short reads. 870 contigs were left after error correction with RACON and Pilon software.

### 2.3 Chromosome assembly using Hi-C

Bowtie2(Langmead & Salzberg, 2012) with the default parameters was used to map the clean reads to the *A. indica*. HiC-Pro v2.11.1(Servant et al., 2015) was used to map the Hi-C sequencing reads to the assembled draft genome and detect the valid contacts. Then we used ALLHIC v0.9.12(Zhang, Zhang, Zhao, Ming, & Tang, 2019) to cluster contigs into chromosome-scale scaffolds based on the relationships among valid contacts.

### 2.4 Assessment of genomic integrity

The draft genome sequence of *A. indica* (GCA_000439995.3) was downloaded from NCBI as a reference. The accuracy and integrity of the genome assembly was evaluated using BUSCO v3.0.2, based on the OrthoDB (http://cegg.unige.ch/orthodb) database. The transcriptomic NGS short reads from 5 tissues of *A.indica* (SRR12709585, SRR12709584, SRR12709583, SRR12709582, and SRR12709581) (Wang, Wang, & Huo, 2020) were mapped against the assemblies using Hisat2(Kim, Langmead, & Salzberg, 2015) with default parameters. The genomic NGS short reads were also mapped to the assemblies using Bowtie2. Finally, the collinearity analysis between our assembly and GCA_000439995.3 was performed with Minimap2 (https://github.com/galaxyproject/tools-iuc/tree/master/tools/minimap2) and dotPlotly (https://github.com/tpoorten/dotPlotly).

### 2.5 Repetitive elements

We identified repetitive elements through both RepeatModeler v1.0.8 (Price, Jones, & Pevzner, 2005) and RepeatMasker v4.0.7(Tarailo-Graovac & Chen, 2009). The LTRs of *A. indica* were identified by using LTRharvest v1.6.1(Ellinghaus, Kurtz, & Willhoeft, 2008) and LTR_Finder v1.05(Z. Xu & Wang, 2007). LTR_retriever v2.8.7 (Ou & Jiang, 2018) was used to integrate the results of LTRharvest and LTR_Finder. RepeatModeler employed RECON v1.08 and RepeatScout v1.0.5 to predict interspersed repeats and then combined the repeat sequences from LTR-retriever with the repeat sequences from RepeatModeler to be the local repeat library. To recover the repeats in the *A. indica* genome, a homology-based repeat search was conducted by using RepeatMasker with the *ab initio* repeat database and Repbase (https://www.girinst.org/repbase/).

### 2.6 Noncoding RNAs

Noncoding RNAs were detected through searching against various RNA libraries. Reliable tRNA positions were searched via tRNAscan-SE v1.3.1(Lowe & Eddy, 1997). Small nuclear RNAs (snRNAs) and microRNAs (miRNAs) were searched by using INFERNAL v1.1(Nawrocki & Eddy, 2013) against the Rfam(Griffiths-Jones et al., 2005) database.

### 2.7 Gene prediction

Homology annotation was performed using genomes of three representative species, including *Citrus sinensis*(Q. Xu et al., 2013), *Theobroma cacao*(Argout et al., 2011), and *Acer yangbiense*(J. Yang et al., 2019). The TBLASTN software(Camacho et al., 2009) was used to align the protein sequences of these species to *A. indica* genome sequence, with an E-value ≤1e-5. The exact gene structures were predicted using GeneWise 2.2.0(Birney, Clamp, & Durbin, 2004) according to the TBLASTN results. We used Cufflinks v2.2.1(Trapnell et al., 2012) to preliminarily identify gene structures based on the RNA-seq data. *ab initio* annotation was performed using Augustus v3.2.2(Stanke, Steinkamp, Waack, & Morgenstern, 2004) and SNAP(Korf, 2004) with the repeat-masked genome sequences. All genes predicted from the three annotation procedures were integrated with MAKER(Holt & Yandell, 2011) software.

### 2.8 Functional annotation

The protein sequences of the consensus gene set were aligned to four protein databases, including NR (https://www.ncbi.nlm.nih.gov/protein/), InterPro (https://www.ebi.ac.uk/interpro/), Swiss-Prot (http://www.uniprot.org), and eggNOG(Powell et al., 2012), for predicted gene annotation. The physically clustered specialized metabolic pathway genes were identified by the PlantiSMASH analytical pipeline(Kautsar, Suarez Duran, Blin, Osbourn, & Medema, 2017). Plant disease resistance (R) genes were predicted by the Disease Resistance Analysis and Gene Ontology (DRAGO) pipeline(Osuna-Cruz et al., 2018).

### 2.9 Phylogenetic analysis and expansion/contraction of gene families

The genome of *A. indica* and 13 other plants were selected for phylogenetic analysis. All-vs.-all BLASTP(Altschul et al., 1997) searching results with an E-value ≤1e-5 were grouped into orthologous and paralogous clusters using OrthoFinder v2.3.7(Emms & Kelly, 2019). Multiple sequence alignments of all single-copy orthologous genes were performed by using MUSCLE(Edgar, 2004). Divergence time between species was estimated using MCMCtree, which was incorporated in the PAML v4.8 package(Z. Yang, 1997). CAFÉ v3.1(De Bie, Cristianini, Demuth, & Hahn, 2006) was used to measure the expansion/contraction of orthologous gene families.

### 2.10 Genome duplication analysis

MCScan v0.8(Tang et al., 2008) package with default parameters was used for the detection of syntenic blocks, defined as regions with more than 5 collinear genes. The YN00 NG model was used to calculate the synonymous substitution rate (*Ks*) between the syntenic homologous gene pairs by PAML(Z. Yang, 1997). The WGD events of each species were estimated based on the *Ks* distributions. The gene pairs with the median *Ks*< 0.05 were defined as the retained genes from the recent segmental duplication. According to the formula *T*=*Ks*/2*r*, the *Ks* values were converted to divergence times, where *T* is divergence time and *r* is the neutral substitution rate (*r* = 3.39 × 10^−9^). The paralog analysis in *A. indica* genome were performed using reciprocal best hits (RBH) from all-vs-all BLASTp searches using *A. indica* protein sequences. RBHs are defined as reciprocal best BLASTp matches with e-value threshold of 1e-5, c-score threshold of 0.3(Guo et al., 2018).

### 2.11 Identification and phylogenetic analysis of TPS and CYP family members

Genomes were aligned using HMMER 3.0 search with an E-value 1e-5 against the Pfam-A database (02-May-2020) locally. PF01397 (Terpene synthase, N-terminal domain) and PF03936 (Terpene synthase family, metal binding domain) domains were used to identify the members of the TPS gene family. The collection used for phylogenetic analysis consisted of 403 putative TPSs from *A. indica* and other 13 plants and six reported TPSs belonged to TPS- a (AAX16121.1), b (AAQ16588.1), c (AAD04292.1), e (Q39548.1), f (Q93YV0.1), and g (ADD81294.1) subfamilies(Y. Kumar, Khan, Rastogi, & Shasany, 2018; Zhou, Shamala, Yi, Yan, & Wei, 2020). PF00067 (Cytochrome P450) was used to identify the members of the CYP gene family. Putative CYPs were screened by amino acid length (450 < length < 600) to perform phylogenetic analysis. Protein sequences were aligned using ClustalX in MEGAX using default sets(S. Kumar, Stecher, Li, Knyaz, & Tamura, 2018). The Neighbor-Joining (NJ) trees were constructed based on the alignment of TPS and CYP protein sequences using MEGAX software with 100 bootstrap replicates, respectively. The identification of *A. indica*-specific TPS and CYP genes was based on the phylogenetic analysis using other 13 plant genome as the outgroup and a cutoff of 55% identity, which indicated separate subfamily assignment(X. Liu et al., 2018; Tu et al., 2020).

### 2.12 Molecular docking analysis

Modelling of CYPs was performed using Phyre2. The 3DLigandSite was used to characterize the potential active site of binding sites. Molecular docking was performed with Autodock 4.0. Four triterpenoids (azadirone, nimbin, nimbolide, and tirucalla-7,24-dien-3β-ol) were selected as potential substrates for CYPs(Wang et al., 2020).

## 3. Results

### 3.1 Genome sequencing and assembly

To obtain a chromosome-level assembly of *A. indica*, the genome was sequenced using a combination of Illumina, PacBio, and Hi-C methods, and assembled by a hierarchical approach. A total of 110 Gb (providing 188 × genome coverage) Illumina paired-end short reads were produced and the heterozygosity ratio was estimated to be 0.896%. Based on the 21-mer depth distribution of the Illumina short reads, the genome size was estimated to be 165 Mb (Figure S1).

We also generated 126 Gb of raw PacBio sequencing reads from the single-molecule real-time (SMRT) sequencing platform, reaching 256 × coverage of the *A. indica* genome (Figure S2, Table S1). The total size of the reads assembled from the post-correction genome was 281,629,231 bp with a GC content of 32.2%, consisting of 870 contigs. The contig N50 was 6,039,544 bp, and the longest contig was 15,111,501 bp.

We further conducted the Hi-C sequencing to scaffold the preliminary assemblies and enhance the assembled contiguity at the chromosome level. In total, the Hi-C sequencing generated approximately 40.48 Gb clean reads. 94.5% reads from Hi-C sequencing were mapped to the assembled contigs, of which 26.1% were unique mapped read pairs (Table S2). The verified read pairs were selected after considering the map position and orientation of the unique mapped read pairs. Then, according to the contiguity information between Hi-C read pairs, ALLHIC software was used to cluster, order, and orient the previous assemblies for chromosome-level scaffolding (Figure 1a). A total of 70 scaffolds were obtained after Hi-C sequencing reads assist chromosome assemble, of which 14 scaffolds formed chromosomes (Figure 1b, Table S3). The final size of the *A. indica* genome assembly was 281 Mb, and the scaffold N50 was 19 Mb (Table 1).

**Table 1.** Statistics of the *A. indica* genome assembly.

| Feature | value |
| --- | --- |
| Genome size (Mb) | 281 |
| Genome GC% | 32.2 |
| N50(Mb) | 19 |
| Gene number | 25,767 |
| Average gene length(bp) | 2,837 |
| Exon no. per gene | 5.4 |
| Exon number | 138,941 |
| Average exon length(bp) | 231 |
| Total exon length(bp) | 32,191,037 |

**Figure 1.**
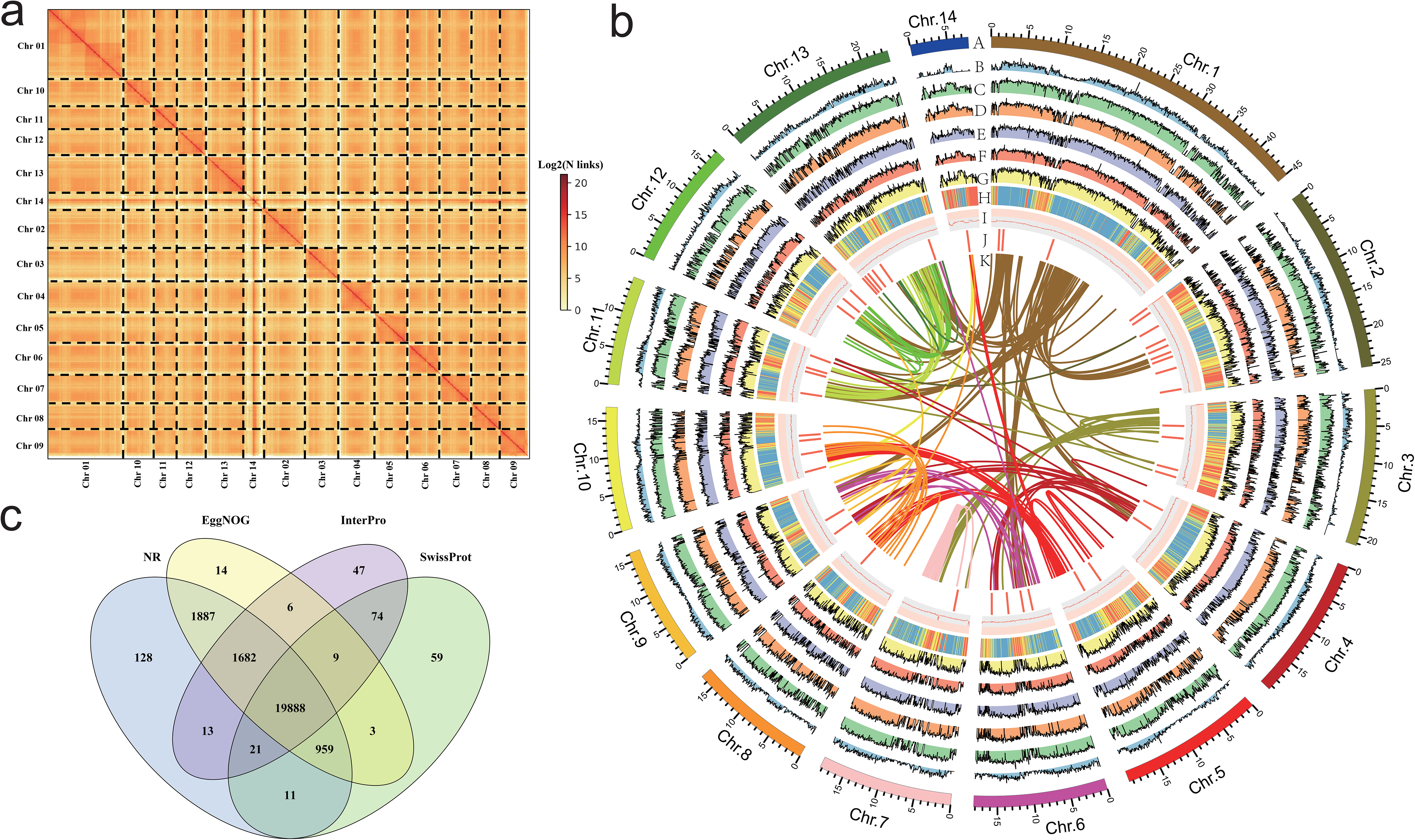
Genome features of genome assembly of *A. indica.* **a** Hi-C contact data mapped on the *A. indica* genome showing genome-wide all-by-all interactions. **b** The landscape of genome assembly and annotation of *A. indica*. (A) Circular representation of the pseudomolecules. (B) The distribution of gene density with densities calculated in 500 kb windows. (C-G) Expression of *A. indica* genes (from outside to inside tracks: stem, root, leaf, fruit and flower). (H-I) The distribution of repeat density and GC density with densities calculated in 500 kb windows. (J) Locations of genes mapped to secondary metabolism. (K) Syntenic blocks. **c** Venn diagram showing the gene function annotation results in NR, InterPro, SwissProt and EggNOG.

### 3.2 Evaluation of the genome assembly

The quality of the assembly was assessed and compared with the reference genome sequence of *A. indica* from NCBI (GCA_000439995.3) (Figure S3, Table 2). The Benchmarking Universal Single-Copy Orthologs (BUSCO)(Simao, Waterhouse, Ioannidis, Kriventseva, & Zdobnov, 2015) analysis was used to evaluate the integrity of the genome. The BUSCO assessment showed that the completeness of the assembled genome of *A. indica* was 91.7%, which was much higher than that of the reference genome (Figure S3a, Table2, Table S4). The Illumina short reads were also used to assess the integrity of the genome. The transcriptomic Illumina sequencing short reads were mapped to the two assemblies by Hisat2(Kim et al., 2015), and approximately 92.76% and 87.49% of the reads were mapped to our assembly and GCA_000439995.3, respectively. By using Bowtie2(Langmead & Salzberg, 2012) software, the genomic Illumina sequencing short reads were also mapped to the assemblies. About 99.29% and 97.09% of the Illumina short reads could map to our assembly and GCA_000439995.3, respectively (Figure S3b). Finally, collinearity analysis revealed good collinearity between our assembly and GCA_000439995.3 (Figure S3c).

**Table 2.** Comparison of the *A. indica* genome assembly versions.

| Feature | This study | Krishnan. et al.10 | GCA_000439995.3 |
| --- | --- | --- | --- |
| Sequence technology | Illumina+PacBio+Hi-C | Illumina+PacBio | Illumina |
| Assembly level | chromosome | scaffold | contig |
| Genome size (Mb) | 281 | 216 | 264 |
| Genome GC% | 32.2 | 31.9 | 32.0 |
| Number of scaffolds | 70 | 25,560 | 126,142 |
| Scaffold N50(bp) | 19,542,739 | 2,629,187 | 3,491 |
| Number of contigs | 870 | 48,555 | 142,701 |
| Contig N50(bp) | 6,039,544 | 25,406 | 3,310 |
| BUSCO | 91.7% | 91.4% | 79.9% |

### 3.3 Gene prediction and genome annotation

Gene models were generated by a combination of reference plant protein homology support, transcriptome data, and *ab initio* gene prediction. All gene models were merged with MAKER(Holt & Yandell, 2011), resulting in a total of 25,767 protein-coding genes with an average sequence length of 2,837 bp. On average, each predicted gene contained 5.4 exons with a mean sequence length of 231 bp (Table 1). In addition, 3,856 noncoding RNAs, including 1,381 rRNAs, 1,204 tRNAs, 173 microRNAs (miRNAs), and 1,098 small nuclear RNAs (snRNAs) were identified (Table S5). We also identified 40.99% of the assembled sequences as repetitive sequences, which was higher than that of the reported genomes(Krishnan et al., 2016; Kuravadi et al., 2015). The majority of the repeats were long terminal repeats (LTRs), constituting 16.88% of the genome. Unclassified elements, DNA elements, and long interspersed nuclear elements (LINEs), accounted for 14.28%, 6.54%, and 1.08% of the genome, respectively (Table S6).

To further evaluate the functional validity of the predicted genes, Diamond, BLASTP, InterProScan and eggnog-mapper were utilized by searching the Nr, SwissProt, InterPro and EggNOG databases (Figure 1c). Overall, 24,801 genes (96.2%) were functionally assigned. 95.4% and 81.6% of these genes found homologies and annotated proteins in the Nr and SwissProt databases, respectively. 84.3% of the genes were detected with conserved protein domains using InterProScan. In addition, 47.4% of the genes were categorized by Kyoto Encyclopedia of Genes and Genomes (KEGG) pathway(Moriya, Itoh, Okuda, Yoshizawa, & Kanehisa, 2007) (Table S7).

### 3.4 Phylogenetic analysis

To investigate the genetic diversity and evolutionary history of *A. indica* genome, a gene family clustering analysis with the *A. indica* genome and 13 other representative plant species was performed. These selected species included two plants in the *Sapindales* order (*Acer yangbiense* and *Citrus sinensis*), eight plants in the eudicot clade (*Arabidopsis thaliana*, *Theobroma cacao*, *Gossypium raimondii*, *Carica papaya*, *Vitis vinifera*, *Cucumis sativus*, *Fragaria vesca*, *Prunus persica*, and *Solanum lycopersicum*), and two outgroup species (*Brachypodium distachyon* and *Amborella trichopoda*).

OrthoFinder(Emms & Kelly, 2019) was used to construct a phylogenetic tree with 1,338 single-copy orthologous genes among 14 species, which showed that *A. indica* is most closely related to *C. sinensis* (Figure 2). Further analysis showed that 36 gene families were specific to *A. indica* (Table S8). Enrichment analysis showed that these specific genes were mostly involved in “binding”, “catalytic activity”, “metabolic process”, “cellular process” and “membrane” (Table S9). With the divergence time between *P. persica* and *F. vesca* as a calibration point (with the corrected time obtained from TimeTree(S. Kumar, Stecher, Suleski, & Hedges, 2017)), the divergence time among these species were also estimated. *A. indica* and *C. sinensis* diverged from a common ancestor ~57 Mya (Figure 2). To better understand the genetic basis of *A. indica*, the expansion and contraction of gene families were investigated. 997 gene families were expanded in *A. indica*, while 293 gene families were contracted from the *A. indica* genome. Compared with *C. sinensis*, which has 369 expanded gene families and 682 contracted gene families, *A. indica* has expanded more gene families. GO and KEGG analysis of the expanded and contracted gene families were also performed (Figure S4, Table S10, and Table S11). The *A. indica*-specific expanded and contracted gene families might be related to the adaptation to *A. indica*-specific tropical niches. Further researches are required to verify the function of these genes.

**Figure 2.**
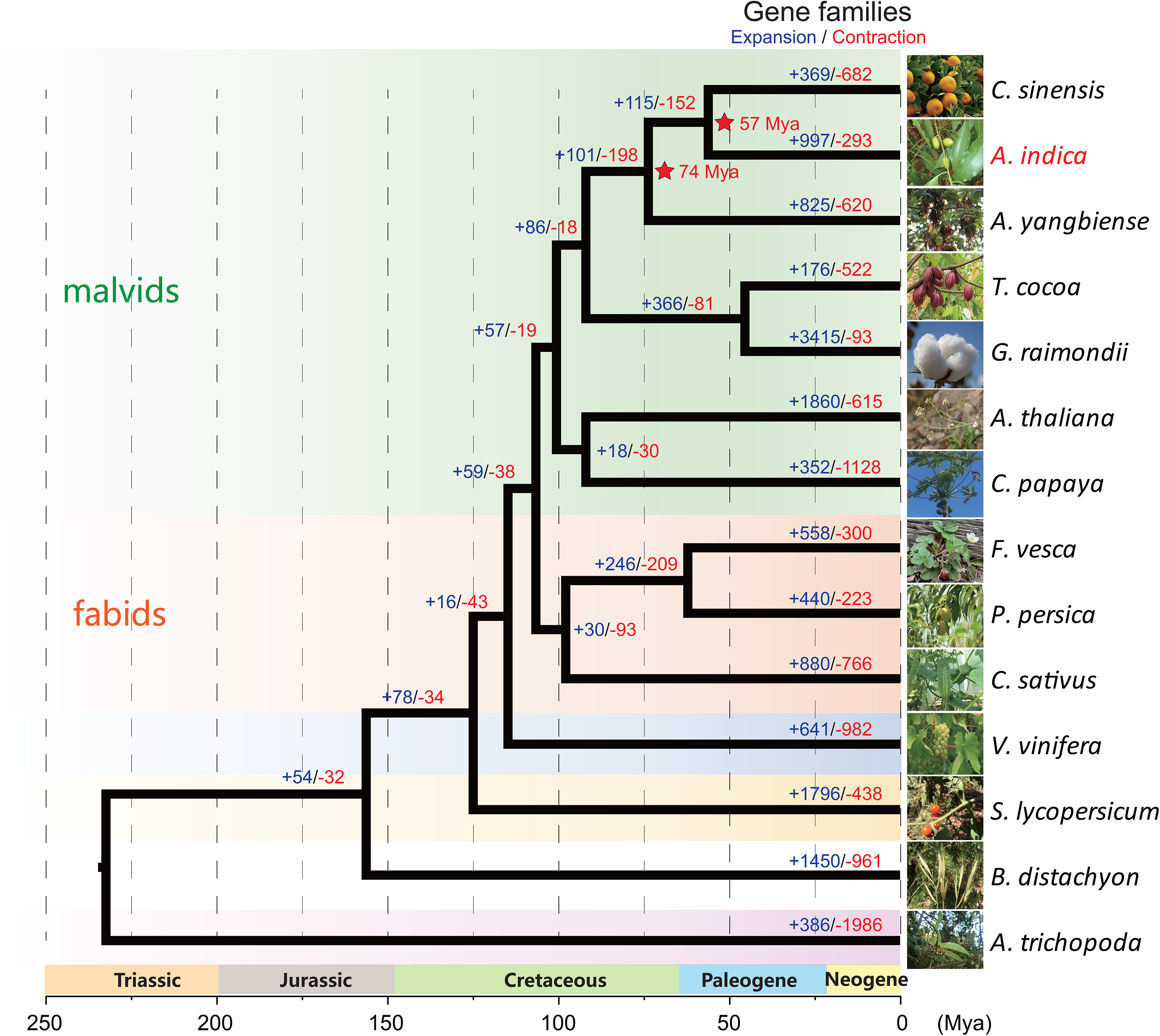
Phylogenetic tree of *A. indica* and 13 other species.

### 3.5 Genome duplication analysis

To investigate genome wide duplications in *A. indica* genome, self-comparison of the *A. indica* genome was performed using MCScan(Tang et al., 2008) (Figure S5). 255 homologous blocks were identified in the intragenomic gene synteny of *A. indica*, containing 2,583 gene pairs. These homologous blocks were distributed across the 14 chromosomes, covering 17.66% of protein-coding genes (4,524/25,767). The synonymous nucleotide substitutions (*Ks*) of the gene pairs peaked at approximately 0.01 and 1.12 (Figure 3a). The first peak at approximately 1.12 indicated the core eudicot γ triplication event (~165 Mya). The second peak at approximately 0.01 indicated a relatively recent duplication event. To distinguish whether this peak represents a whole genome duplication event or background duplications, we performed synteny analysis on *A. indica*, *V. vinifera*, *C. sinensis* and *A. yangbiense* genomes (Figure 3b, Figure S6). Intergenomic collinearity analysis showed 371 homologous blocks containing 16,402 gene pairs and a 3:3 syntenic relationship between *A. indica* and *V. vinifera* (Figure 3b, Figure S7a). Although there were 2:1 syntenic relationship between *A. indica* vs. *C. sinensis* and *A. indica* vs. *A. yangbiense* (Figure S7b, S7c), only 13% and 14% of the *A. indica* gene models in syntenic blocks, respectively, were present as two copies. Meanwhile, we did not identify large *C. sinensis* and *A. yangbiense* segments that have two syntenic copies in *A. indica* by the synteny dot plot of *A. indica* vs. *C. sinensis* and *A. indica* vs. *A. yangbiense* (Figure S6). Our analysis indicated that *A. indica* didn’t experience additional WGD after the γ event, but a recent small-scale segmental duplication(Q. Xu et al., 2013; J. Yang et al., 2019). The calculation of *Ks* for *A. indica* vs. *C. sinensis* indicated that this recent segmental duplication event occurred approximately 1.5 Mya after the divergence from *C. sinensis.* Furthermore, we also performed paralog analysis in *A. indica* genome using reciprocal best hits (RBH) from primary protein sequences by all-vs-all BLASTp matches. We detected 6,496 RBH paralogous gene pairs in the *A. indica* genome, and the RBH paralog *Ks* distribution shows a *Ks* peak at around 0.01 (Figure S8). That this RBH *Ks* peak is close to the syntelog *Ks* peak also indicates *A. indica* has a recent segmental duplication mixed with gene duplication.

**Figure 3.**
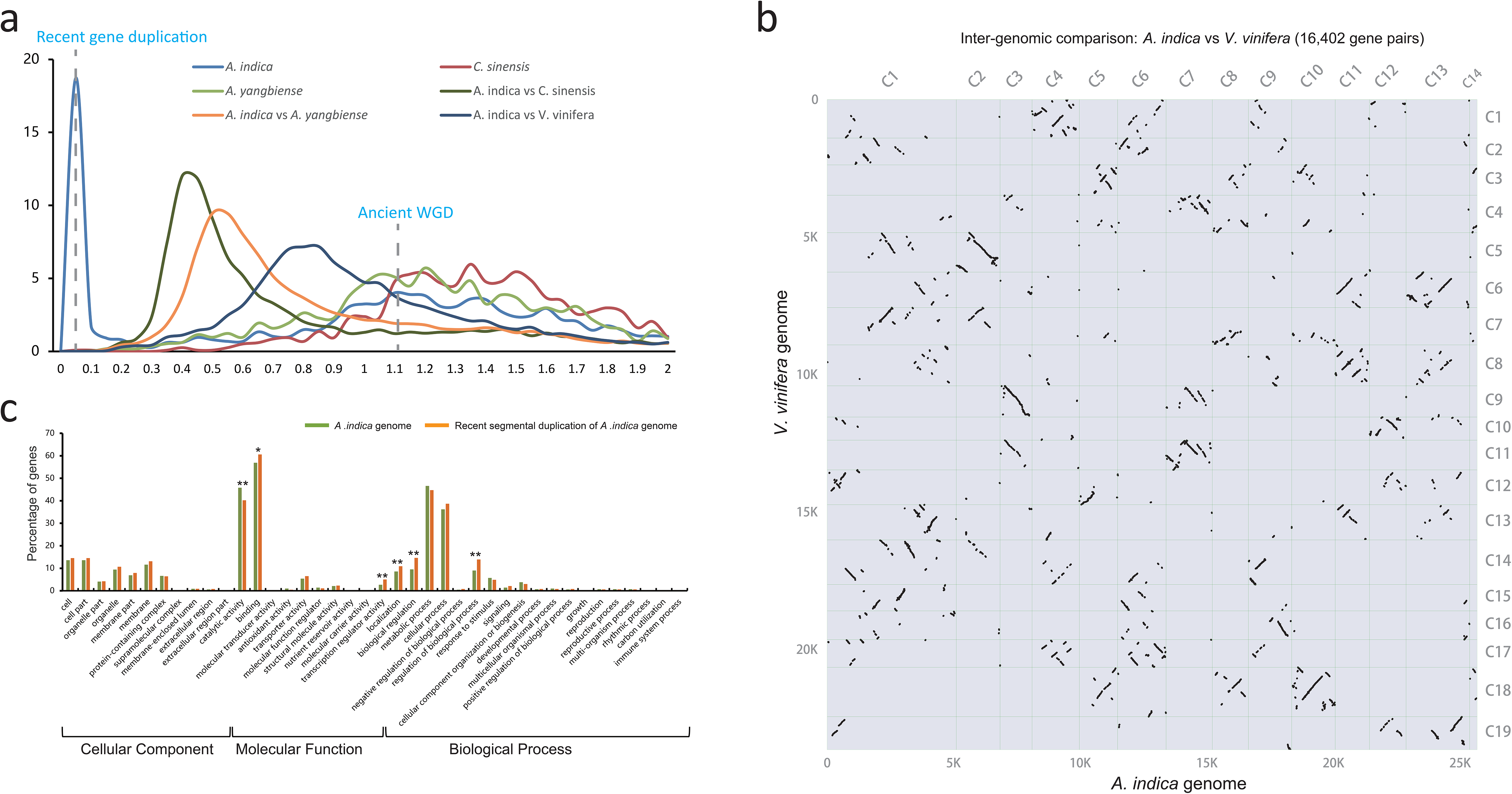
Genome evolution of *A. indica*. **a** *Ks* distribution (The gene pairs located in syntenic blocks) between syntenic genes within the *A. indica* genome or between genomes. **b** Inter-genomic syntenic analysis between *A. indica* genome and *V. vinifera* genome. **c** Enrichment of the *A. indica* genome (green) and genes retained after the recent gene duplication (orange). *, Pearson Chi-Square test P value < 0.05; **, Pearson Chi-Square test P value < 0.01.

Generally, gene duplication events vary the genomic architecture, including genome size, genome density, gene content, and gene expression. In this study, we defined the gene pairs that located in syntenic blocks with the median *Ks* < 0.05 as the retained genes from the recent gene duplication. 1,539 homologous gene pairs retained after the recent gene duplication. GO analysis revealed that these gene pairs were involved in binding, catalytic activity, metabolic process, and cellular process (Figure 3c). The recent gene duplication may also affect the percentage of genes in many function categories with different contributions. In the *A. indica* genome, the percentage of retained genes from recent gene duplication in “binding”, “transcription regulator activity”, “localization”, “biological regulation”, and “regulation of biological process” was much greater than that of the average genome content (Figure 3c, Figure S9). We further calculated the omega values (*Ka*/*Ks*) for most of the homologous gene pairs. Most of the omega values for the homologous gene pairs were smaller than 1, which indicated that purifying selection may be the predominant action within the retained genes from the recent segmental duplication (Stix, 1992). However, 116 gene pairs were identified that have experienced potential positive selection. GO analysis showed that these genes were mainly enriched in “heterocyclic compound binding”, “organic cyclic compound”, “organic substance metabolic process”, and “primary metabolic process” (Table S12, Figure S10).

### 3.6 Secondary metabolite analysis

Genes encoding some specialized metabolic pathways are found physically clustered in plant genomes(Z. Liu et al., 2020; Nutzmann, Huang, & Osbourn, 2016). We utilized the PlantiSMASH analytical pipeline(Hu et al., 2019) to identify physically clustered specialized metabolic pathway genes. According to the analysis, 50 clusters including 692 genes were identified in the *A. indica* genome (Table S13). The sizes of the identified clusters range from 27.2 kb to 1634.4 kb. 105 (out of 692) clustered genes were contained in the 997 *A. indica*-specific expansion gene families (Table S14). Furthermore, 41(*C. sinensis*), 51 (*A. yangbiense*), 48 (*T. cocoa*), 47 (*G. raimondii*), 45 (*A. thaliana*), 35 (*F. vesca*), 33 (*P. persica*), 30 (*C. sativus*), 46 (*V. vinifera*), 47 (*S. lycopersicum*), and 29 (*B. distachyon*) clusters were detected in other 11 species (Figure 4a). As expected, more terpene-related clusters were identified in the *A. indica* genome than that of other species.

**Figure 4.**
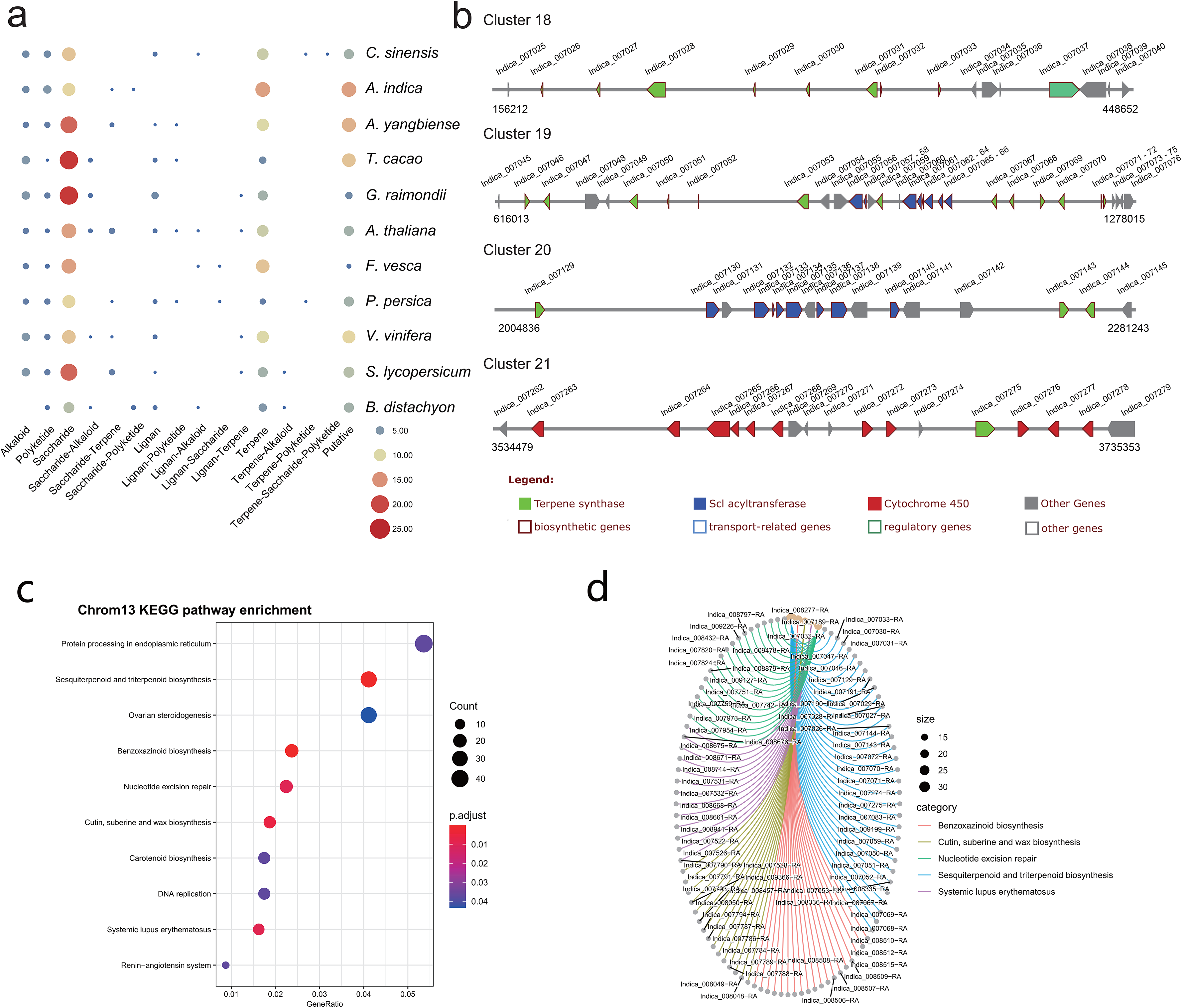
Secondary metabolite analysis. **a** Secondary metabolite analysis of *A. indica* and other 11 species. **b** The organization and architecture of terpene-related gene clusters on chromosome 13. **c** Dotplot of the KEGG pathway enrichment analysis. **d** The correlation between enrichment pathway and genes on chromosome 13.

Azadirachtin is a triterpenoid compound of neem tree, which has effective insecticidal activities against a wide range of insect species, but has very low toxicity to mammals. Terpene synthase (TPS), cytochrome P450 (CYP450), alcohol dehydrogenase (ADH), acyltransferase (ACT), and esterase (EST) were proposed to be involved in biosynthesis of azadirachtin(Wang et al., 2020). In this study, a large number of genes encoding CYP P450s (78), TPSs (58), and ACTs (34) were identified in secondary metabolite biosynthesis gene clusters. Genes encoding ADHs, and ESTs may reside dispersedly in the genome. The terpene-related clusters mainly distributed on chromosome 1, 2, 3, 5, 6, 7, 10, 11, 12, and 13. Four terpene-related clusters (cluster 18-21) covering ~1.4 Mb were distributed on chromosome 13 (Figure 4b). Among the 83 clustered terpene-related genes on chromosome 13, 12 genes were contained in the *A. indica*-specific expanded gene families. These genes are proposed to be potential genes participated in the terpene biosynthesis specific to *A. indica*.

KEGG enrichment analysis was further performed to investigate the function of genes on chromosome 13. The result showed that genes on chromosome 13 were mainly involved in “Protein processing in endoplasmic reticulum”, “Sesquiterpenoid and triterpenoid biosynthesis”, and “Ovarian steroidogenesis” (Figure 4c). According to the cnetplot, 33 genes were correlated with the “Sesquiterpenoid and triterpenoid biosynthesis” pathway (Figure 4d).

### 3.7 Terpene synthase gene family

TPS gene family is characterized by two large domains: PF01397 (Terpene synthase, N-terminal domain) and PF03936 (Terpene synthase family, metal binding domain). To investigate the characteristics and evolution of the TPS gene families, we identified a total of 512 putative TPS genes in *A. indica* and other 13 plant genome. 70 putative TPS genes were identified in *A. indica*; These consisted of 44 AziTPS genes containing both PF01397 and PF03936 domains, nine AziTPS genes containing PF01397 domain, and 17 AziTPS genes containing PF03936 domain. *A. indica* (N=70) contained the most copies of TPSs compared with other plants, followed by *C. sinensis* (N=49) and *A. yangbiense* (N=57) (Figure 5a, Table S15 and Table S16).

**Figure 5.**
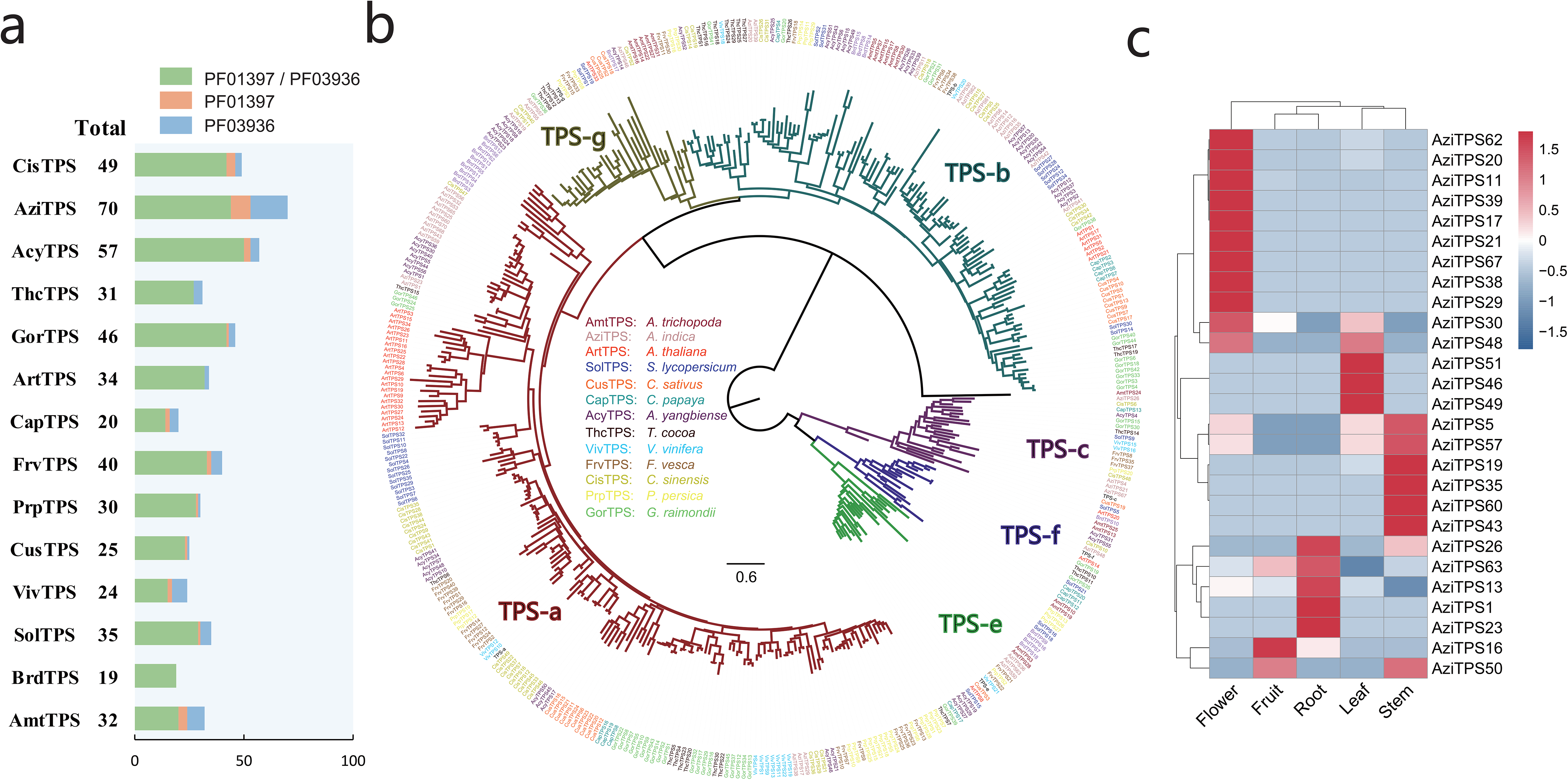
Analysis of TPS gene family in *A. indica*. **a** Statistics of predicted TPS genes in *A. indica* and other 13 plants. **b** Phylogenetic analysis of TPS family members based on the protein sequences. A Neighbor-Joining (NJ) tree was generated from an alignment of 409 TPS protein sequences, comprising 403 putative TPSs from *A. indica* and other 13 plants (the remaining TPS genes were too short for meaningful alignment) and other six reported TPS genes belonged to TPS- a (AAX16121.1), b (AAQ16588.1), c (AAD04292.1), e (Q39548.1), f (Q93YV0.1), and g (ADD81294.1) subfamilies. **c** *A. indica* TPS genes substantially expressed in five tested organs (root, flower, fruit, leaf, and stem).

Phylogenetic analysis was performed using 403 TPSs (the remaining TPS genes were too short for meaningful alignment) from *A. indica* and other 13 plants, including six reported TPS genes belonged to TPS- a, b, c, e, f, and g subfamilies, respectively (Table S17). As shown in Figure 5b, the topology of six subfamilies is similar to that of the previous papers(Ji et al., 2021; Y. Kumar et al., 2018; Zhou et al., 2020). Among the 40 AziTPS used in phylogenetic analysis, 15, 13, 4, 3,1, and 4 AziTPS genes fell in TPS- a, b, c, e, f, and g subfamilies, respectively. TPS-a and -b subfamilies were the main subfamilies in *A. indica*, approximately 37.5% and 32.5% of the total AziTPS genes in phylogenetic analysis. This is in accordance with other plant species, including tea, grape, and Chinese mahogany(F. Chen, Tholl, Bohlmann, & Pichersky, 2011; Ji et al., 2021; Zhou et al., 2020). Furthermore, we identified putative *A. indica*-specific TPSs using phylogenetic analysis and a cutoff of 55% identity, which indicates separate subfamily assignment(X. Liu et al., 2018; Tu et al., 2020). A total of nine *A. indica*-specific TPS genes were identified (Table S18). Interestingly, seven of these specific AziTPSs (Indica_007028, Indica_007047, Indica_007053, Indica_007068, Indica_007070, Indica_007072, and Indica_007143) were located in the terpene-related clusters (cluster 18, 19, and 20) of chromosome 13 (Table S18).

We further investigated the expression pattern of TPS genes in *A. indica*. Transcriptome datasets from five tissues of *A. indica* were obtained from our previous study(Wang et al., 2020) and remapped to the chromosome-level genome assembly in this study. More than 88% of the RNAseq reads were mapped uniquely to the genome assembly across all samples (Table S19). Transcripts of 27 TPS genes were detected in the tested tissues. Most of the detected transcripts exhibited a spatial-specific expression pattern (Figure 5c). Nine, one, four, three, and four genes were exclusively expressed in flower, fruit, root, leaf, and stem, respectively. Seven genes (AziTPS30, −48, −5, −57, −26, −63, and −50) were primarily expressed in one or two tissues.

### 3.8 Cytochrome P450 gene family

The characteristics and evolution of the cytochrome P450 (CYP) gene families were also investigated. In total, 3657 CYP genes were identified from all 14 plant genomes, 355 of these CYP genes were in the *A. indica* genome (Figure 6a, Table S20, and Table S21). Moreover, 157 full length CYP (450 < length < 600) protein sequences of *A. indica* were aligned to construct a phylogenetic tree. As shown in Figure 6b, the phylogenetic tree was divided into two major clades: A type (49%; 77/157) and non-A type (51%; 80/157); and further clustered into nine clan. The Clan 71 is the largest clan and comprises of 49% (77/157) members; 18, 4, 28, and 25 members are classified into Clan72, Clan74, Clan85, and Clan86; remaining Clan51, Clan710, Clan711, and Clan727 are single family clans.

**Figure 6.**
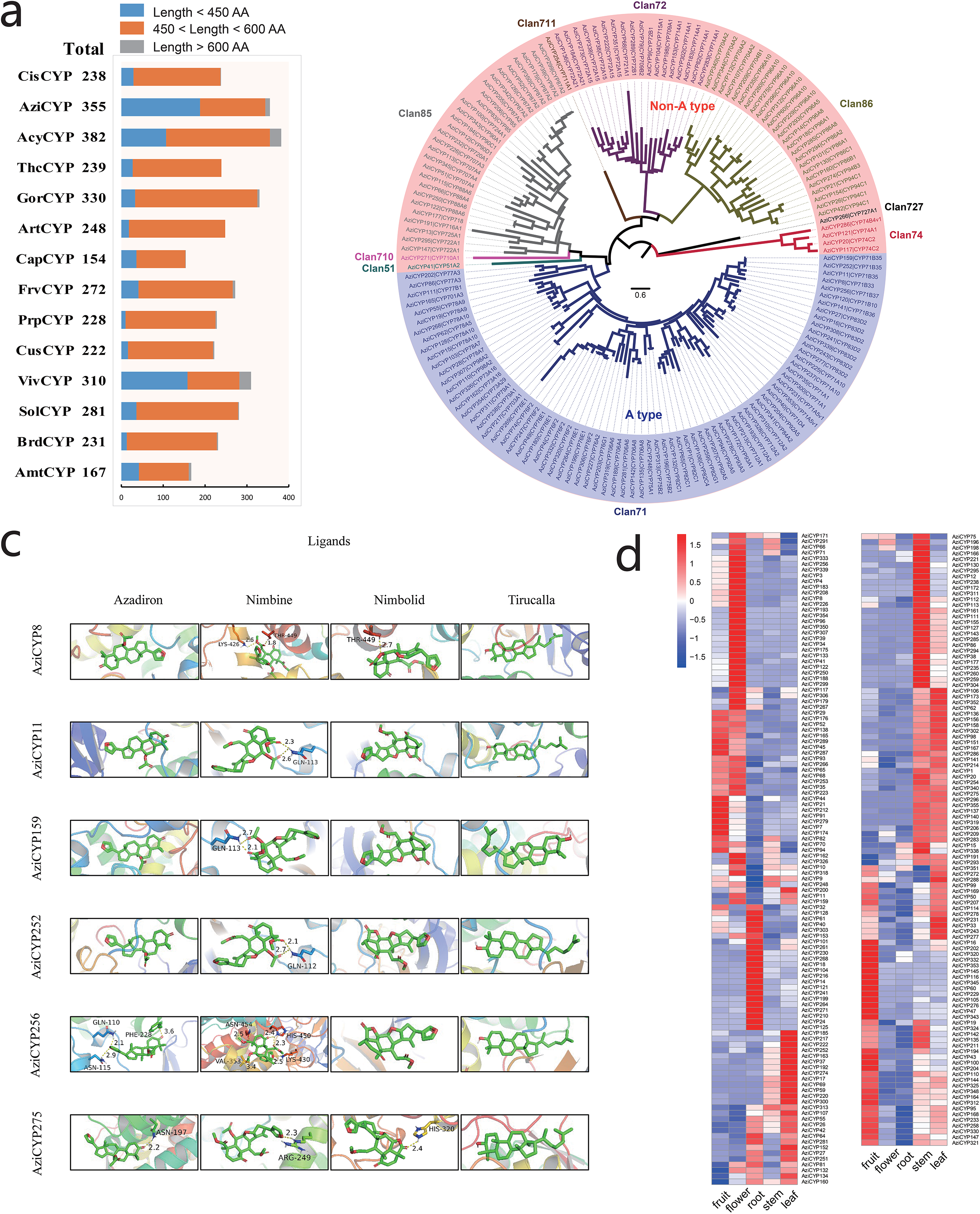
Analysis of P450 gene family in *A. indica*. **a** Statistics of predicted P450 genes in *A. indica* and other 13 plants. **b** Phylogenetic analysis P450 family members in *A. indica* based on the protein sequences. A NJ tree was generated from an alignment of 156 *A. indica* P450 protein sequences (450 < length < 600). The entire family of P450 genes is shown for each clan next to the tree with different color. **c**. Molecular docking analysis of *A. indica*-specific CYPs. Interaction of six CYPs docked regions with four ligands (azadirone, nimbin, nimbolide, and tirucalla-7,24-dien-3β-ol) are shown. **d** *A. indica* CYP genes substantially expressed in five tested organs (fruit, flower, root, stem, and leaf).

In order to identify putative *A. indica*-specific CYP genes, we constructed a phylogenetic tree using amino acid sequence alignment of 2807 (450 < length < 600) CYP genes in *A. indica* and other 13 plants genome with a cutoff of 55% identity(X. Liu et al., 2018; Tu et al., 2020). Six *A. indica-*specific CYP genes were identified (Table S22). Similar to TPS genes, five of these CYP genes (Indica_007272, Indica_007273, Indica_007276, Indica_007277, and Indica_007278) were located in the terpene-related cluster 21 of chromosome 13 (Table S22). These specific-TPSs and CYPs in the terpene-related secondary metabolite biosynthesis gene clusters of chromosome 13 might be involved in the specific biosynthesis of azadirachtin. Molecular docking analysis of *A. indica*-specific CYP genes were further performed (Figure 6c). The active site predication and the details of docking are listed in Table S23 and S24, respectively. The binding energy of AziCYP8, AziCYP11, AziCYP159, AziCYP252, and AziCYP256 was lowest for tirucalla-7,24-dien-3β-ol with −10.2, −9.9, −9.4, −10.8, and −10 kcal/mol without forming hydrogen bond, respectively. Docking of AziCYP275 showed interaction through one hydrogen bond (HIS-320) with the lowest binding energy of −8.3 kcal/mol for nimbolide. Nimbin with AziCYP8 and AziCYP256 and nimbolide with AziCYP256 formed stable complexes through multiple hydrogen bonds.

We also investigated the expression pattern of *A. indica* CYP genes in different tissues (fruit, flower, root, stem, and leaf). Transcripts of 221 CYP genes were detected with different patterns (Figure 6d). There were more high-expressed CYPs in fruit, stem and leaf than flower and root. The high-expressed CYPs in fruit, stem and leaf were 83, 88 and 97, respectively. CYPs with a high-expression in the tissues (fruit and leave) with high azadirachtin A content, are more likely to be involved in azadirachtin biosynthesis. Furthermore, *A. indica*-specific AziCYP256 (Indica_007272) and AziCYP8 (Indica_007273) were highly expressed in fruit and flower.

### 3.9 Resistance genes

Plants have developed a wide range of defense mechanisms to protect themselves against the attack of pathogens in their constant struggle for survival. In general, proteins encoded by resistance (R) genes display modular domain structures. In this study, putative R genes in the *A. indica* genome (1488) and other 13 species were identified (Table S25). In the *A. indica* genome, 238 R genes may exert their disease resistance function as cytoplasmic protein through canonical resistance domains, such as the nucleotide-binding sites (NBSs), the leucine-rich repeat (LRR) and terminal inverted repeat (TIR) domains (Table S26). 167 NBS genes were identified in the *A. indica* genome, which could be divided into five classes according to the conserved domains: N, CN, CNL, NL, and TNL. The majority were N type which contained only the NB-ARC domain. In comparison with other genomes in malvids, most of the NBS genes in the *A. indica* genome were underrepresented relative to other Sapindales genomes (*C. sinensis* and *A. yangbiense*) and Malvales genomes (*T. cacao* and *G. raimondii*), but overrepresented relative to other Brassicales genome (*A. thaliana* and *C. papaya*). In addition, 447 genes were classified as transmembrane receptors, including 221 receptor-like kinases (RLK), and 226 receptor-like proteins (RLP). 721 putative kinases were also identified in the *A. indica* genome.

## 4. Discussion

*A. indica* is a valuable plant species given its economic and pharmaceutical significance(Stix, 1992). A high-quality reference genome is essential for the genetic and genomic studies of *A. indica*. However, molecular-level studies on this species are limited. Here, we assembled the first chromosome-scale genome of *A. indica* by a combination of Illumina, PacBio, and Hi-C technology. The size of the genome assembly is approximately 268M, with a scaffold N50 value of 19 Mb. The N50 of our assembled genome is much higher than that of the previous published draft genomes(Krishnan et al., 2016; Krishnan et al., 2012; Kuravadi et al., 2015). The *A. indica* genome shows a high level of heterozygosity (0.896%) and repeat content (40.99%), rendering substantial challenges for its assembly(Nowak et al., 2015). Hi-C technology has been broadly available for many complex species(J. D. Chen et al., 2020). In this study, Hi-C technology facilitated the completeness and accuracy of a chromosome-level genome assembly for *A. indica*. The improvement of BUSCO evaluation shows that our assembly represents a better template for gene annotation than the reference sequence. Considering that the genome is highly heterozygous and repetitive, the present version represents a high-quality genome assembly. The obtained genome is also the second chromosome-level genome of the Meliaceae family, which will pave the way for further genetic and genomic studies of the Meliaceae family.

Gene duplication is an important evolutionary force that provides abundant raw materials for genetic novelty, morphological diversity and speciation (Qiao et al., 2018). In this study, no WGD event occurred in *A. indica* genome after the ancient γ event shared by the eudicot genome. However, recent gene duplication events mixed with small-scale segmental duplication may occur in multiple genes in *A. indica*. This may also explain the fact that *A. indica* had more expanded gene families than *C. sinensis*. Considering that the recent gene duplication event occurred during a period of rapid cooling with transient intervals of CO_2_ reduction(Pearson & Palmer, 2000), it is likely that the recent gene duplication event helped increase the environmental robustness and the potential for specific adaptation of *A. indica*. All these results are highly benefit for in-depth investigation of the survival and diversification history the of Meliaceae family.

Limonoids are natural triterpenoid products made by plants of the Meliaceae family. They are known for their insecticidal activity and potential pharmaceutical properties. *A. indica* is known as the reservoir of azadirachtin, the most famous limonoid insecticide. Secondary metabolite analysis revealed that *A. indica* contained more terpene-related clusters than that of the other 11 species. 83 (out of 247) clustered terpene-related genes were located on chromosome 13. The KEGG pathway enrichment analysis revealed that 33 genes were correlated with the “Sesquiterpenoid and triterpenoid biosynthesis” pathway. These results indicated that chromosome 13 may play an unneglectable role for the terpene biosynthesis of *A. indica*.

TPS and CYP gene family is responsible for the biosynthesis of terpenoids in plants. 70 TPS genes were identified in *A. indica*, which is much more than that of the other 13 species. This is consistent with the result of Chinese mahogany (*Toona sinensis*), the first chromosome-level genome assembly of the Meliaceae family(Ji et al., 2021). 355 CYP genes were identified in *A. indica*, six of which were *A. indica* specific CYPs. The expansion of terpene-related gene clusters, TPSs and CYPs, may promote the formation of terpenoids in *A. indica*. Notably, most of the identified *A. indica* -specific TPSs and CYPs were located in the terpene-related clusters on chromosome 13, indicating that these regions were likely to be involved in azadirachtin biosynthesis. This study provided the first chromosome-level genome of *A. indica*, and a genomic perspective for the synthesis and evolution of azadirachtin.

## Supporting information

Table S10 KEGG pathways of expanded families in A. indica

Table S11 KEGG pathway of contracted families in A. indica

Table S12 Positively selected genes from the recent gene duplication

Table S13 The gene list of the secondary metabolite clusters in the A. indica genome

Table S14 The gene list of the secondary metabolite clusters from the A. indica specific expanded gene families

Table S15 Identification of the TPS genes in A. indica genome

Table S16 The statistics of the TPS genes identified in the A. indica genome and 13 other plant genomes

Table S17 The statistics of the TPS genes in phylogenetic tree

Table S18 Identification of the A. indica-specific TPS genes

Table S19 Transcriptome data quality control and mapping results

Table S20 Identification of the P450 genes in A. indica genome

Table S21 The statistics of the P450 genes identified in the A. indica genome and 13 other plant genomes

Table S22 Identification of the A. indica-specific P450 genes

Table S23 Potential active site identified on the A. indica-specific CYP450s

Table S24 Details of molecular docking of the A. indica-specific CYPs with probable ligands

Table S25 The statistics of the resistance genes identified in the A. indica genome and 13 other plant genomes

Table S26 Resistance genes in the A. indica genome

## Acknowledgements

This study was funded by the National Key R&D Program of China (2017YFD0201400), the Fundamental Research Funds for the Central Universities, and the General Program of National Natural Science Foundation of China (31970622).

## Data availability

Raw data from this study were deposited in the NCBI SRA (Sequence Read Archive) database under the Bioproject ID: PRJNA645650. The genome sequence data (Illumina, PacBio, and Hi-C data) are available under accession numbers SRR12315383, SRR12321691, and SRR12321285. The assembled genome was submitted to DDBJ/ENA/GenBank with accession number JAGQDM000000000. The metadata is available at https://dataview.ncbi.nlm.nih.gov/object/PRJNA645650?reviewer=137oln8itc913iv8p2i8fbrn2b.

## Author contributions

Y. D. and W. S. designed the project and wrote the draft manuscript. W. S. participated in the genome assembly and annotation. Y. D. and Z. Y. contributed to the genome evolution analysis, gene family analysis, and resistance gene identification. S. W. and J. L. performed the molecular docking analysis. N. W., H. J., J. Q. and Y.-X. H. revised the manuscript. The final manuscript has been read and approved by all authors.

## Supplementary information

**Figure S1**. The distribution of A. indica 21-mers. The frequency of each 21-mer was calculated based on the filtered paired-end reads from libraries with short inserts (150bp). Two peaks were observed at x and x, respectively.

**Figure S2**. Read length distribution of PacBio sequencing data.

**Figure S3.** Comparison of the *A. indica* genome assembly with the reference assembly. **a** Comparison of the BUSCO evaluation between our assembly and the reference assembly. Red, missing (M); Yellow, fragmented (F); Dark blue, complete (C) and duplicated (D); Wathet, complete (C) and single-copy (S). **b** Comparison of the transcriptomic Illumina short reads mapping (left) and genomic Illumina short reads mapping (right) between our assembly and the reference assembly. **c** Collinearity between our assembly and the reference assembly.

**Figure S4**. Enriched GO terms of *A. indica* expanded (a) or contracted (b) genes.

**Figure S5**. Intra-genomic syntenic analysis of *A. indica*.

**Figure S6.** Inter-genomic syntenic analysis of *A. indica*-vs-*C. sinensis* (a) and *A. indica*-vs-*A. yangbiense* (b).

**Figure S7**. Summary of the syntenic analysis of *A. indica*-vs-*vinifera* (a), *A. indica*-vs-*C. sinensis* (b) and *A. indica*-vs-*A. yangbiense* (c).

**Figure S8.** *Ks* distribution for *A. indica* RBH (reciprocal best hit) paralogs.

**Figure S9**. The comparison in gene numbers and percentages of significantly different GO terms between recent segmental duplication and whole genome.

**Figure S10**. Enrichment analysis of the positively selected genes from the recent segmental duplication.

**Table S1**. The statistics of sequencing raw data from the Pacific Biosciences RS II sequencing platform.

**Table S2**. Statistics of Hi-C data and assessment.

**Table S3**. Chromosome length by Hi-C assembly.

**Table S4**. Quality assessment of the assembled genome of *A. indica* using BUSCOs.

**Table S5**. Statistics on the annotation of non-coding RNA of the *A. indica* genome.

**Table S6**. Repeats in the *A. indica* genome assembly.

**Table S7**. Summary of the functional annotation in *A. indica* genome.

**Table S8**. Summary of gene family clustering in *A. indica*.

**Table S9**. The list of *A. indica* specific genes.

**Table S10**. KEGG pathways of expanded families in *A. indica*.

**Table S11**. KEGG pathway of contracted families in *A. indica*.

**Table S12**. Positively selected genes from the recent gene duplication.

**Table S13**. The gene list of the secondary metabolite clusters in the A. indica genome.

**Table S14**. The gene list of the secondary metabolite clusters from the *A. indica* specific expanded gene families.

**Table S15**. Identification of the TPS genes in *A. indica* genome.

**Table S16**. The statistics of the TPS genes identified in the *A. indica* genome and 13 other plant genomes.

**Table S17**. The statistics of the TPS genes in phylogenetic tree.

**Table S18**. Identification of the *A. indica*-specific TPS genes.

**Table S19**. Transcriptome data quality control and mapping results.

**Table S20**. Identification of the P450 genes in *A. indica* genome.

**Table S21**. The statistics of the P450 genes identified in the *A. indica* genome and 13 other plant genomes.

**Table S22**. Identification of the *A. indica*-specific P450 genes.

**Table S23**. Potential active site identified on the *A. indica*-specific CYP450s.

**Table S24**. Details of molecular docking of the *A. indica*-specific CYPs with probable ligands.

**Table S25**. The statistics of the resistance genes identified in the *A. indica* genome and 13 other plant genomes.

**Table S26**. Resistance genes in the *A. indica* genome.

## Genome Assemblies

Genome assemblies are not expected to satisfy all of these areas, but rather it is necessary to generally demonstrate high quality genomes that will be strong resources for broader studies for the community.

It is now policy at Molecular Ecology Resources for genome assemblies and all data associated with an assembly to be made available for the editor and reviewers at time of submission. This includes FASTA assembly files and raw read data such as RNAseq data. Please detail how editors and reviewers can access this data in the Cover Letter.

- Were whole genome shotgun libraries sequenced at high coverage for the target species (provide in terms of “X coverage “)? 188×coverage.
- Did the study generate a pan-genome assembly from several individual samples? No.
- Were ‘long-read’ libraries sequenced and included in the genome assembly (specify type and coverage)? PacBio sequence was used and reached 256 × coverage of the *A. indica* genome.
- What are the basic assembly statistics: genome size, percent assembled, # contigs, contig N50, # scaffolds, scaffold N50? genome size:281Mb; percent assembled:91.7%; # contigs:870; contig N50:6Mb; # scaffolds:70; scaffold N50:19Mb.
- Was mapping of some form (genetic, physical, optical) incorporated to order scaffolds? No.
- Were scaffolds anchored to chromosome positions, and if so, what proportion of the genome is anchored to chromosomes? Yes, after Canu software assemble, 870 contigs were obtained. A total of 70 scaffolds were obtained after Hi-C sequencing reads assist chromosome assemble, of which 14 scaffolds formed chromosomes. There are 814 contigs in 14 chromosomes, therefore, 93.6% of the genome was anchored to chromosome.
- Were analyses included that asses the quality of the genome assembly (genomescope, or busco)? BUSCO.
- Were RNAseq libraries sequenced to assemble transcriptomes and annotate genes? Yes.
- Is the genome assembly publicly available through a web-based genome browser? No.
- How is the genome assembly in this manuscript useful for broader research in the field of molecular ecology? This is the first chromosome-level genome assembly of *A. indica*, which will shed light on the biosynthesis of natural products of *A. indica*. This is also the second chromosome-level genome assembly of the Meliaceae, which will facilitate comparative genomic research of the family.
- Please specify the research community that would use this genome resource. Feel free to include names of laboratories. Researchers in plant natural product biosynthesis would use this genome resource, such as Professor Anne Osbourn’s lab in John Innes Centre and Zheyong Xue’s lab in Northeast Forestry Universty.

