## Supplementary information for "Genomic analysis based on chromosome-level genome assembly reveals an expansion of terpene biosynthesis of *Azadirachta indica*"

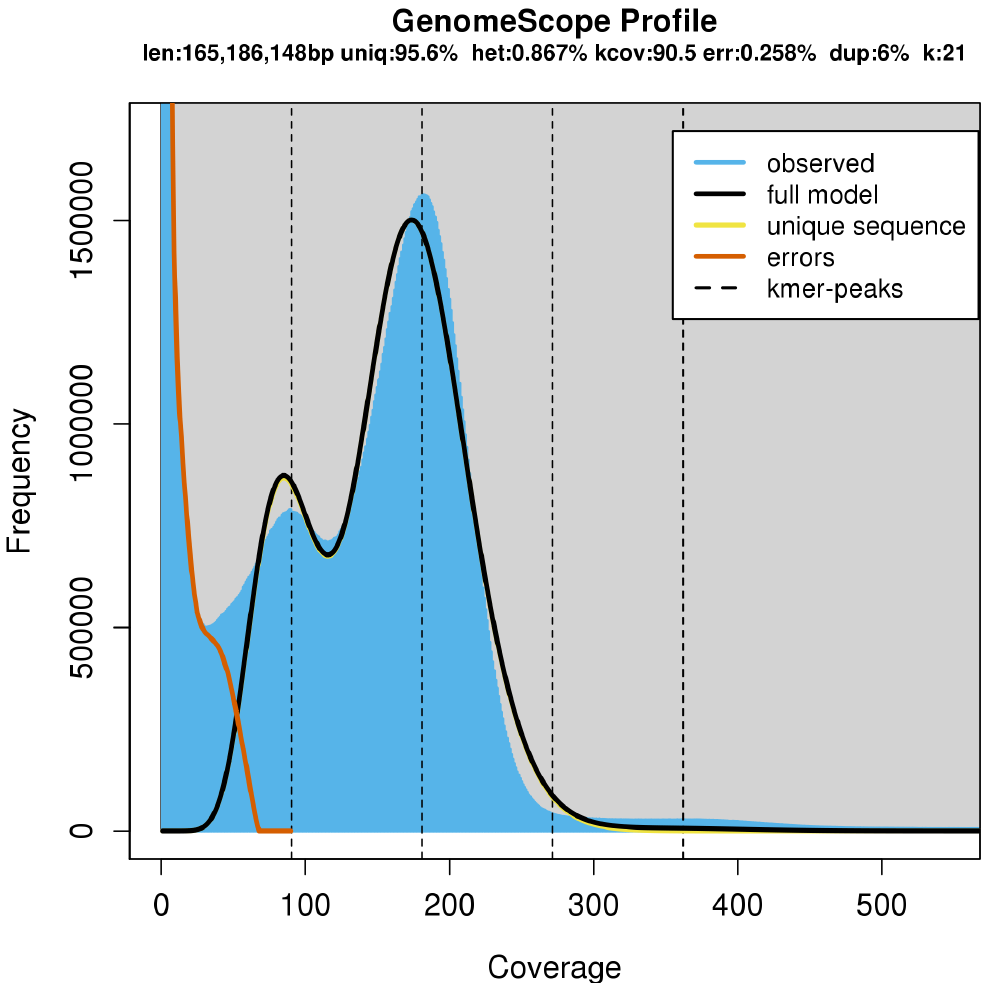

**Figure S1**. The distribution of A. indica 21-mers. The frequency of each 21-mer was calculated based on the filtered paired-end reads from libraries with short inserts (150bp). Two peaks were observed at x and x, respectively.

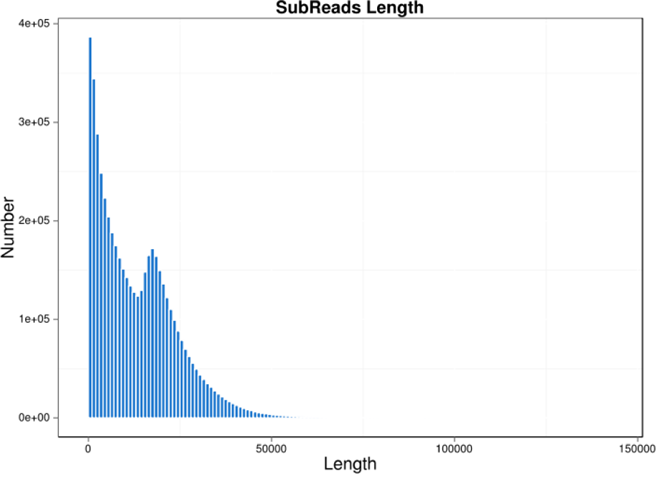

**Figure S2**. Read length distribution of PacBio sequencing data.

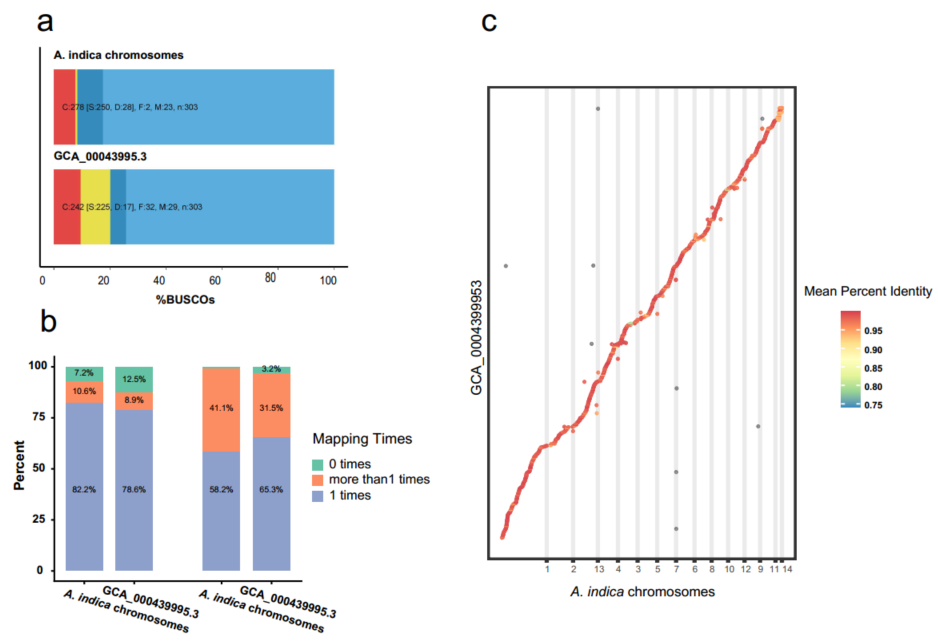

**Figure S3.** Comparison of the *A. indica* genome assembly with the reference assembly. **a** Comparison of the BUSCO evaluation between our assembly and the reference assembly. Red, missing (M); Yellow, fragmented (F); Dark blue, complete (C) and duplicated (D); Wathet, complete (C) and single-copy (S). **b** Comparison of the transcriptomic Illumina short reads mapping (left) and genomic Illumina short reads mapping (right) between our assembly and the reference assembly. **c** Collinearity between our assembly and the reference assembly.

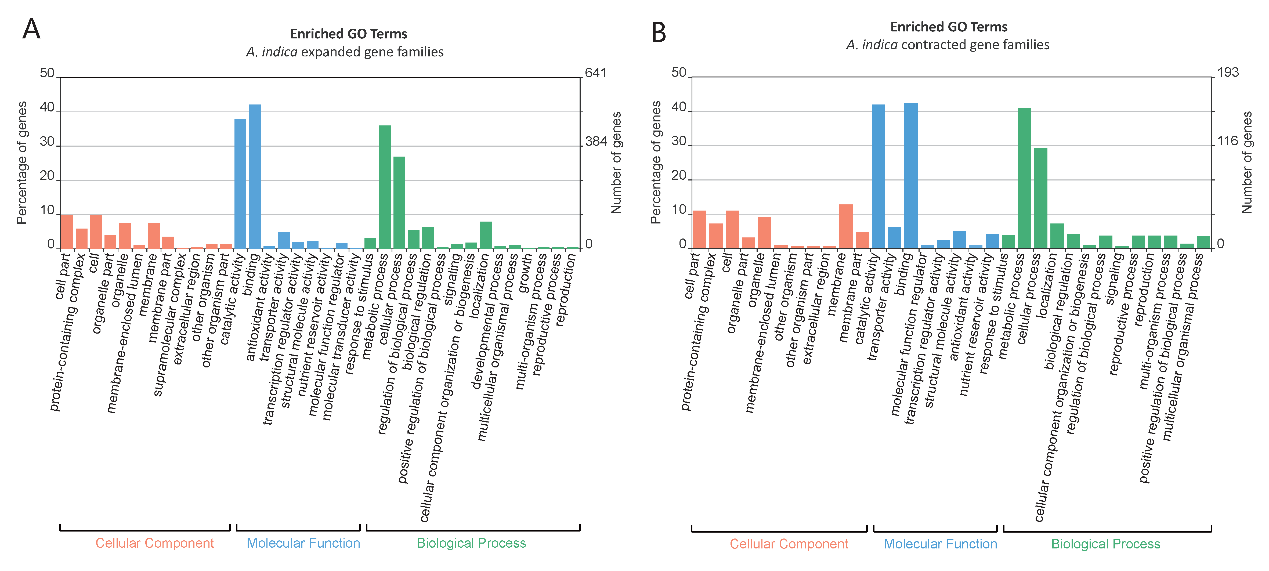

**Figure S4**. Enriched GO terms of *A. indica* expanded (a) or contracted (b) genes.

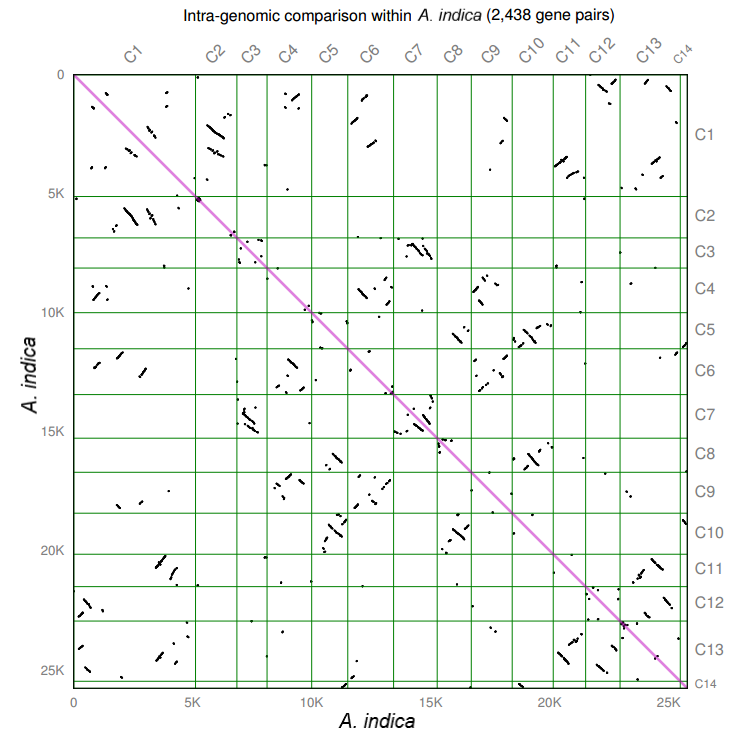

**Figure S5**. Intra-genomic syntenic analysis of *A. indica*.

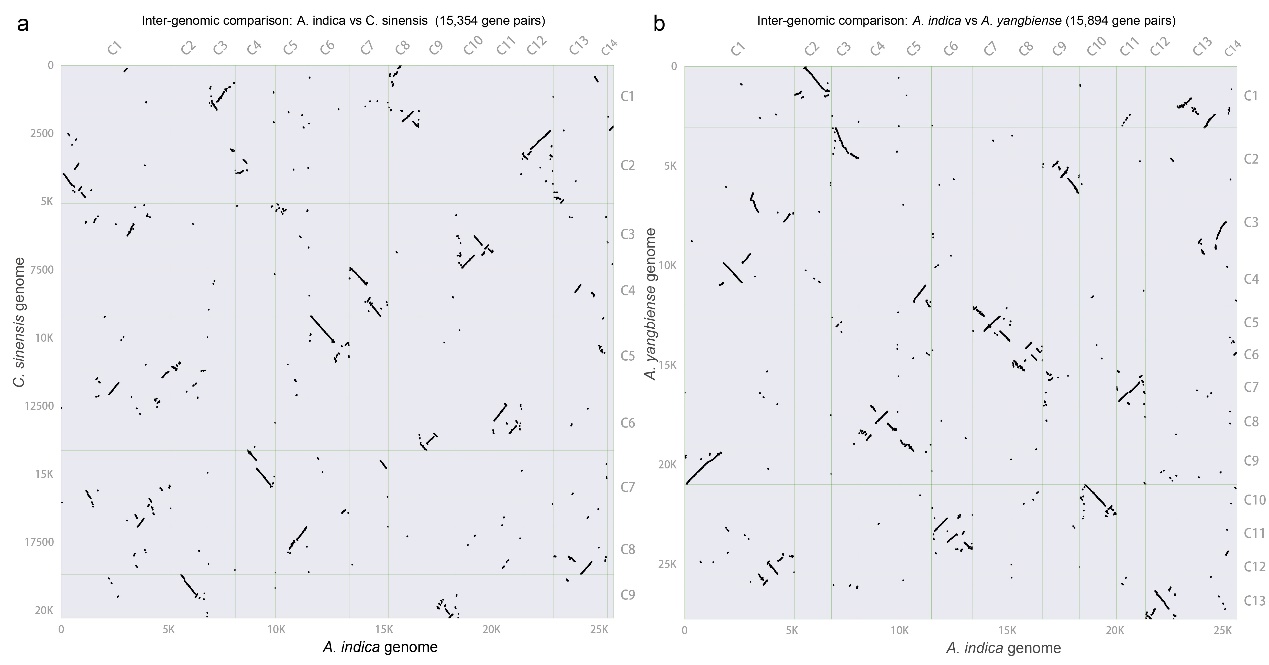

**Figure S6.** Inter-genomic syntenic analysis of *A. indica*-vs-*C. sinensis* (a) and *A. indica*-vs-*A. yangbiense* (b).

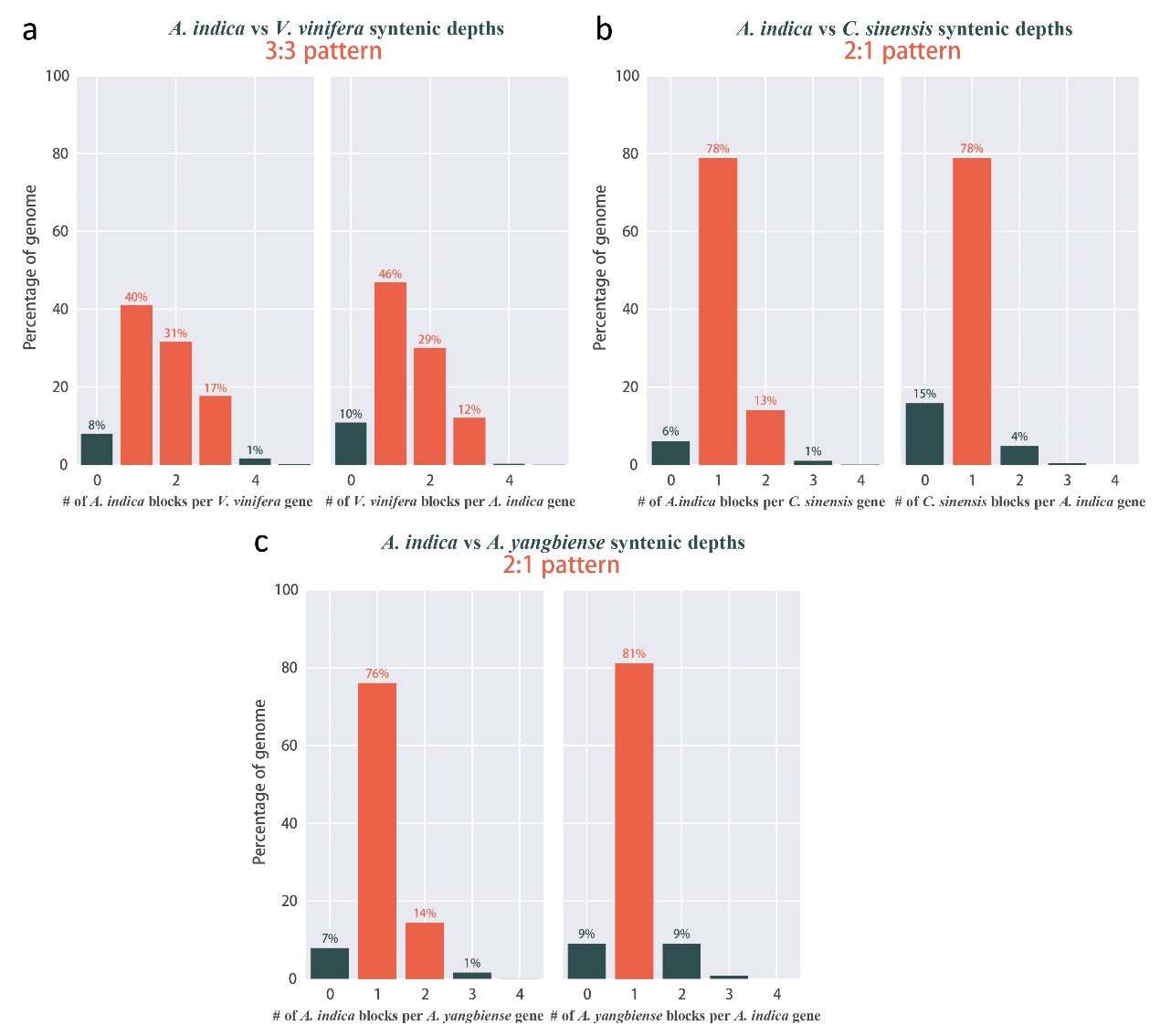

**Figure S7**. Summary of the syntenic analysis of *A. indica*-vs-*vinifera* (a), *A. indica*-vs-*C. sinensis* (b) and *A. indica*-vs-*A. yangbiense* (c).

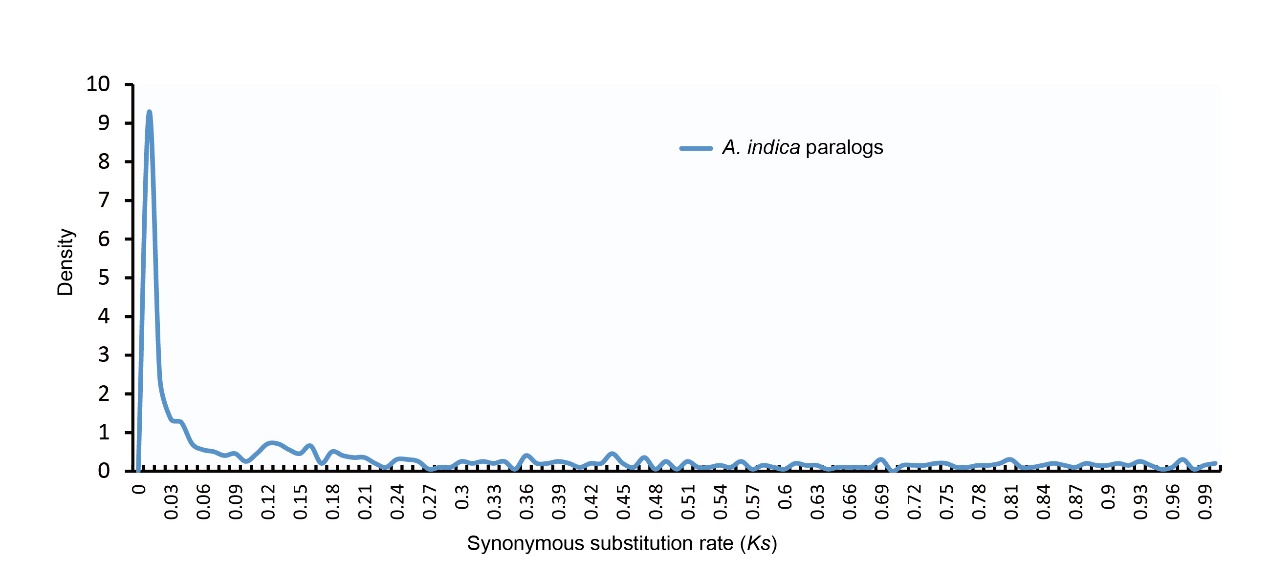

**Figure S8.** *Ks* distribution for *A. indica* RBH (reciprocal best hit) paralogs.

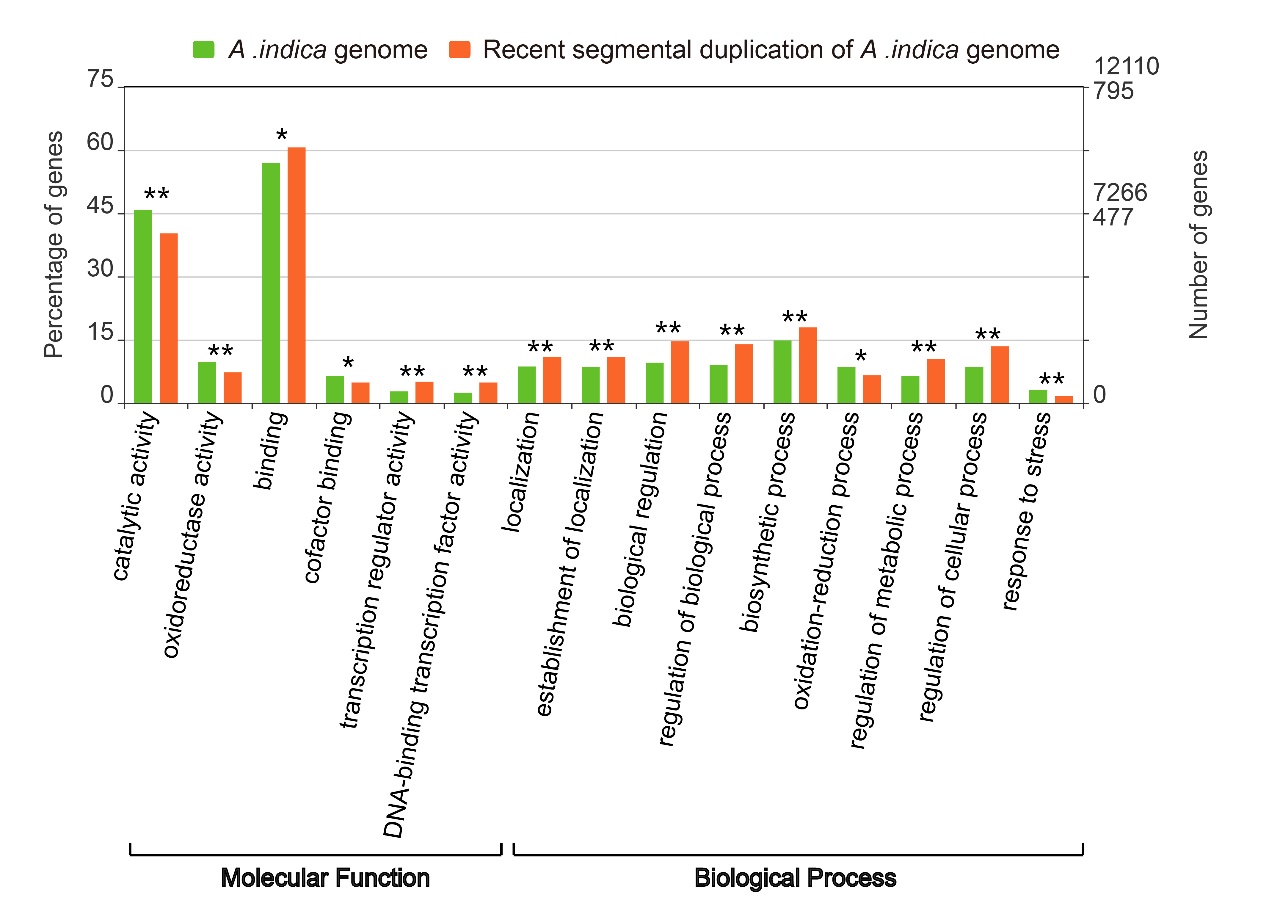

**Figure S9**. The comparison in gene numbers and percentages of significantly different GO terms between recent segmental duplication and whole genome.

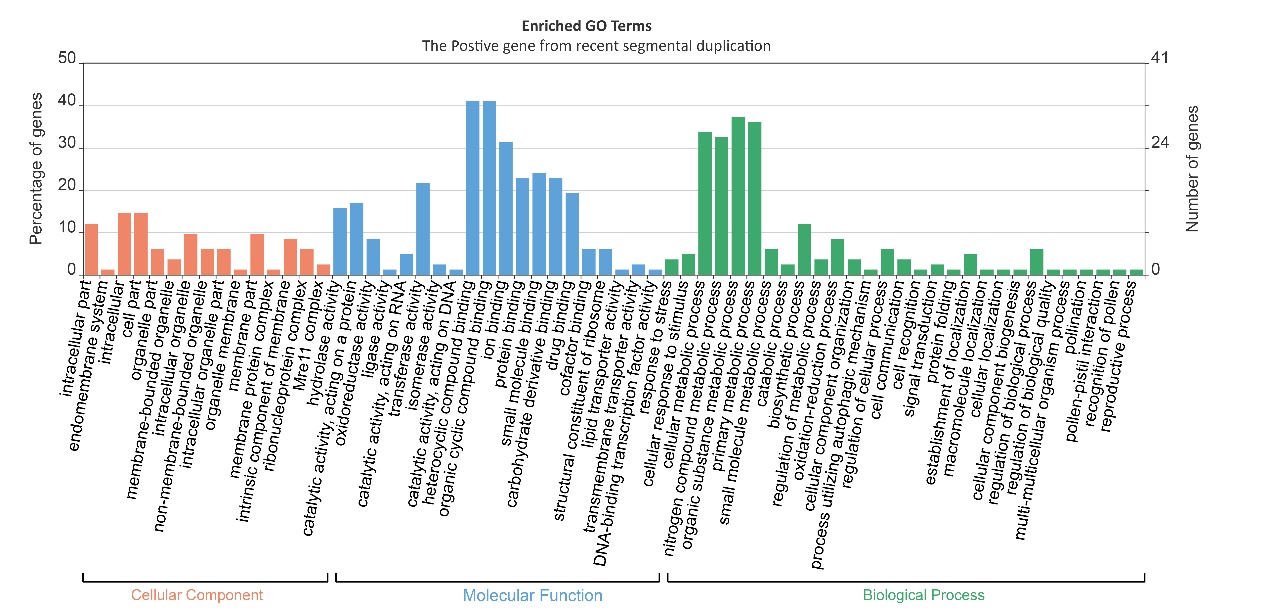

**Figure S10**. Enrichment analysis of the positively selected genes from the recent segmental duplication.

**Table S1**. The statistics of sequencing raw data from the Pacific Biosciences RS II sequencing platform.

| **ID** | **ZMNUM** | **Total Bases**  **(GB)** | **Total reads** | **Average**  **length (bp)** | **Max**  **length (bp)** | **N50 length (bp)** |
| --- | --- | --- | --- | --- | --- | --- |
| 2A | 4341500 | 67.13 | 5072953 | 13233.48 | 142,061 | 20,580 |

**Table S2**. Statistics of Hi-C data and assessment.

| **Statistics of Hi-C data** | | | |
| --- | --- | --- | --- |
| Number of read pairs | Number of bases (bp) | GC content (%) | % ≥ Q30 |
| 209,518,594 | 62,855,578,200 | 38 | 92.34 |
| **Statistics of mapping** | | | |
| Mapping type | | Number of reads | Ratio (%) |
| Total read pairs | | 209,518,594 | 100 |
| Mapped reads | | 197,969,869 | 94.5 |
| Unique mapped read pairs | | 54,648,340 | 26.1 |
| **Statistics of valid Hi-C data** | | | |
| Type | | Number of reads | Ratio (%) |
| Valid interaction pairs | | 42,815,207 | 100 |
| Dangling end pairs | | 9,973,827 | 23.3 |
| Re-ligation pairs | | 1,438,953 | 3.4 |
| Self-cycle pairs | | 412,795 | 0.9 |
| Dumped pairs | | 7558 | 0.018 |

**Table S3**. Chromosome length by Hi-C assembly.

| **Chromosome ID** | **Scaffold Num** | **Length(bp)** |
| --- | --- | --- |
| Chrom. 01 | 68 | 45,097,400 |
| Chrom. 02 | 100 | 25,348,148 |
| Chrom. 03 | 90 | 21,293,100 |
| Chrom. 04 | 59 | 19,556,420 |
| Chrom. 05 | 61 | 19,542,739 |
| Chrom. 06 | 40 | 18,209,783 |
| Chrom. 07 | 46 | 18,270,782 |
| Chrom. 08 | 50 | 16,959,782 |
| Chrom. 09 | 32 | 16,187,903 |
| Chrom. 10 | 59 | 16,873,511 |
| Chrom. 11 | 40 | 14,805,454 |
| Chrom. 12 | 57 | 17,142,395 |
| Chrom. 13 | 54 | 23,763,320 |
| Chrom. 14 | 58 | 7,901,508 |
| total | 814 | 280,952,245 |

**Table S4**. Quality assessment of the assembled genome of *A. indica* using BUSCOs.

| **Type** | **Number** | **Percent (%)** |
| --- | --- | --- |
| Complete BUSCOs (C) | 278 | 91.7 |
| Complete and single-copy BUSCOs (S) | 250 | 82.5 |
| Complete and duplicated BUSCOs (D) | 28 | 9.2 |
| Fragmented BUSCOs (F) | 2 | 0.7 |
| Missing BUSCOs (M) | 23 | 7.6 |
| Total BUSCO groups searched | 303 | 100 |

**Table S5**. Statistics on the annotation of non-coding RNA of the *A. indica* genome.

| **Type** |  | **Copy** | **Total length (bp)** | **% of genome** |
| --- | --- | --- | --- | --- |
| miRNA |  | 173 | 21342 | 0.007596 |
| tRNA |  | 1204 | 90808 | 0.032321 |
| rRNA |  | 1381 | 1895631 | 0.674716 |
| snRNA | CD-box | 408 | 41730 | 0.014853 |
|  | HACA-box | 44 | 5653 | 0.002012 |
|  | splicing | 97 | 14494 | 0.005158 |

**Table S6**. Repeats in the *A. indica* genome assembly.

| **Types** | **No. of copies** | **Length (bp)** | **Coverage of genome (%)** |
| --- | --- | --- | --- |
| DNA transposon | 58968 | 18378069 | 6.54 |
| LINE | 8035 | 3034159 | 1.08 |
| Long terminal repeat | 66790 | 47418507 | 16.88 |
| SINE | 5 | 394 | 0.00 |
| Other∗ | 2925 | 6226012 | 2.21 |
| Unknown | 130734 | 40113882 | 14.28 |
| Total | 267457 | 115171023 | 40.99 |

∗Other includes microsatellites, simple repeats, and low-complexity sequences.

**Table S7**. Summary of the functional annotation in *A. indica* genome.

| **Annotation database** | **Annotated number** | **Percentage (%)** |
| --- | --- | --- |
| Nr | 24,589 | 95.4% |
| SwissProt | 21,024 | 81.6% |
| EggNOG | 24,448 | 94.8% |
| COG | 22,789 | 88.4% |
| InterPro | 21,740 | 84.3% |
| GO | 16,170 | 62.7% |
| KEGG Pathway | 12,126 | 47.4% |
| Total | 24,801 | 96.2% |

**Table S8**. Summary of gene family clustering in *A. indica*.

| **Species** | **Total**  **genes** | **Genes in families** | **Family** | **Unclustered genes** | **Species-specific families** | **Genes per family** |
| --- | --- | --- | --- | --- | --- | --- |
| *A. indica* | 25657 | 22384 | 13398 | 3273 | 36 | 1.67 |
| *A. yangbiense* | 27760 | 24057 | 12931 | 3703 | 187 | 1.86 |
| *C. sinensis* | 20286 | 19367 | 12000 | 919 | 23 | 1.61 |
| *A. thaliana* | 27444 | 22584 | 12286 | 4860 | 347 | 1.84 |
| *T. cacao* | 21257 | 20734 | 13324 | 523 | 17 | 1.56 |
| *G. raimondii* | 35177 | 31903 | 13446 | 3274 | 399 | 2.37 |
| *C. papaya* | 18003 | 16840 | 12404 | 1163 | 19 | 1.35 |
| *V. vinifera* | 23647 | 18177 | 11952 | 5470 | 72 | 1.52 |
| *C. sativus* | 19521 | 18580 | 12174 | 941 | 62 | 1.53 |
| *F. vesca* | 24056 | 22166 | 13083 | 1890 | 91 | 1.69 |
| *P. persica* | 22988 | 21826 | 13186 | 1162 | 58 | 1.66 |
| *S. lycopersicum* | 25157 | 23040 | 12532 | 2117 | 210 | 1.84 |
| *B. distachyon* | 25447 | 20039 | 11586 | 5408 | 670 | 1.73 |
| *A. trichopoda* | 17099 | 15775 | 12042 | 1324 | 178 | 1.31 |

**Table S9**. The list of *A. indica* specific genes.

|  | **GO annotation** | | | **KEGG annotation** | |
| --- | --- | --- | --- | --- | --- |
| **Protein ID** | **Cellular Component** | **Molecular Function** | **Biological Process** | **KO number** | **Definition** |
| Indica_007398-RA |  | GO:0005524\|ATP binding; GO:0004672\|protein kinase activity; | GO:0006468\|protein phosphorylation; | K08829 | MAK; male germ cell-associated kinase [EC:2.7.11.22] |
| Indica_007413-RA |  | GO:0005524\|ATP binding; GO:0004672\|protein kinase activity; | GO:0006468\|protein phosphorylation; | K08829 | MAK; male germ cell-associated kinase [EC:2.7.11.22] |
| Indica_007414-RA |  | GO:0005524\|ATP binding; GO:0004672\|protein kinase activity; | GO:0006468\|protein phosphorylation; | K08829 | MAK; male germ cell-associated kinase [EC:2.7.11.22] |
| Indica_007416-RA |  | GO:0005524\|ATP binding; GO:0004672\|protein kinase activity; | GO:0006468\|protein phosphorylation; | K08829 | MAK; male germ cell-associated kinase [EC:2.7.11.22] |
| Indica_007418-RA |  | GO:0005524\|ATP binding; GO:0004672\|protein kinase activity; | GO:0006468\|protein phosphorylation; | K08829 | MAK; male germ cell-associated kinase [EC:2.7.11.22] |
| Indica_001271-RA |  | GO:0003824\|catalytic activity; |  |  |  |
| Indica_001274-RA |  | GO:0003824\|catalytic activity; |  |  |  |
| Indica_001279-RA |  | GO:0003824\|catalytic activity; |  |  |  |
| Indica_001282-RA |  | GO:0003824\|catalytic activity; |  |  |  |
| Indica_026282-RA |  |  |  |  |  |
| Indica_026294-RA | GO:0016020\|membrane; | | GO:0055114\|oxidation-reduction process; |  |  |
| Indica_026300-RA | GO:0016020\|membrane; | | GO:0055114\|oxidation-reduction process; |  |  |
| Indica_026387-RA | GO:0016020\|membrane; | | GO:0055114\|oxidation-reduction process; |  |  |
| Indica_013831-RA |  | GO:0016787\|hydrolase activity; |  |  |  |
| Indica_013832-RA |  | GO:0016787\|hydrolase activity; |  |  |  |
| Indica_013838-RA |  | GO:0016787\|hydrolase activity; |  |  |  |
| Indica_023874-RA |  | GO:0005515\|protein binding; |  |  |  |
| Indica_023876-RA |  | GO:0005515\|protein binding; |  |  |  |
| Indica_024678-RA |  | GO:0005515\|protein binding; |  |  |  |
| Indica_026283-RA |  |  |  |  |  |
| Indica_026289-RA |  |  |  |  |  |
| Indica_026299-RA |  |  |  |  |  |
| Indica_000067-RA |  | GO:0003676\|nucleic acid binding; |  |  |  |
| Indica_001559-RA |  | GO:0003676\|nucleic acid binding; |  |  |  |
| Indica_021414-RA |  | GO:0005524\|ATP binding; GO:0004672\|protein kinase activity; | GO:0006468\|protein phosphorylation; |  |  |
| Indica_021590-RA |  | GO:0005524\|ATP binding; GO:0004672\|protein kinase activity; | GO:0006468\|protein phosphorylation; |  |  |
| Indica_025821-RA |  |  |  |  |  |
| Indica_025823-RA |  | GO:0008194\|UDP-glycosyltransferase activity; | |  |  |
| Indica_011978-RA |  |  |  | K13343 | PEX14; peroxin-14 |
| Indica_026073-RA |  |  |  | K13343 | PEX14; peroxin-14 |
| Indica_009679-RA |  | GO:0043531\|ADP binding; |  |  |  |
| Indica_010481-RA |  | GO:0043531\|ADP binding; |  |  |  |
| Indica_018790-RA |  |  |  |  |  |
| Indica_019629-RA |  |  |  |  |  |
| Indica_016399-RA |  | GO:0005515\|protein binding; |  |  |  |
| Indica_017175-RA |  | GO:0005515\|protein binding; |  |  |  |
